# Efficient ancestry and mutation simulation with msprime 1.0

**DOI:** 10.1101/2021.08.31.457499

**Authors:** Franz Baumdicker, Gertjan Bisschop, Daniel Goldstein, Graham Gower, Aaron P. Ragsdale, Georgia Tsambos, Sha Zhu, Bjarki Eldon, E. Castedo Ellerman, Jared G. Galloway, Ariella L. Gladstein, Gregor Gorjanc, Bing Guo, Ben Jeffery, Warren W. Kretzschmar, Konrad Lohse, Michael Matschiner, Dominic Nelson, Nathaniel S. Pope, Consuelo D. Quinto-Cortés, Murillo F. Rodrigues, Kumar Saunack, Thibaut Sellinger, Kevin Thornton, Hugo van Kemenade, Anthony W. Wohns, Yan Wong, Simon Gravel, Andrew D. Kern, Jere Koskela, Peter L. Ralph, Jerome Kelleher

## Abstract

Stochastic simulation is a key tool in population genetics, since the models involved are often analytically intractable and simulation is usually the only way of obtaining ground-truth data to evaluate inferences. Because of this necessity, a large number of specialised simulation programs have been developed, each filling a particular niche, but with largely overlapping functionality and a substantial duplication of effort. Here, we introduce `msprime` version 1.0, which efficiently implements ancestry and mutation simulations based on the succinct tree sequence data structure and `tskit` library. We summarise `msprime`’s many features, and show that its performance is excellent, often many times faster and more memory efficient than specialised alternatives. These high-performance features have been thoroughly tested and validated, and built using a collaborative, open source development model, which reduces duplication of effort and promotes software quality via community engagement.

## Introduction

The coalescent process (Kingman, 1982a,b; Hudson, 1983b; Tajima, 1983) models the ancestry of a set of sampled genomes, providing a mathematical description of the genealogical tree that relates the samples to one another. It has proved to be a powerful model, and is now central to population genetics (Hudson, 1990; Hein et al., 2004; Wakeley, 2008). The coalescent is an efficient framework for population genetic simulation, because it allows us to simulate the genetic ancestry for a sample from an idealised population model, without explicitly representing the population in memory or stepping through the generations. Indeed, Hudson (1983b) independently derived the coalescent *in order to* efficiently simulate data, and used these simulations to characterise an analytically intractable distribution. This inherent efficiency, and the great utility of simulations for a wide range of purposes, has led to dozens of different tools being developed over the decades (Carvajal-Rodríguez, 2008; Liu et al., 2008; Arenas, 2012; Yuan et al., 2012; Hoban et al., 2012; Yang et al., 2014; Peng et al., 2015).

Two technological developments of recent years, however, pose major challenges to most existing simulation methods. Firstly, fourth-generation sequencing technologies have made complete chromosome-level assemblies possible (Miga et al., 2020), and high quality assemblies are now available for many species. Thus, modelling genetic variation data as a series of unlinked non-recombining loci is no longer a reasonable approximation, and we must fully account for recombination. However, while a genealogical tree relating *n* samples in the single-locus coalescent can be simulated in *O*(*n*) time (Hudson, 1990), the coalescent with recombination is far more complex, and programs such as Hudson’s classical `ms` (Hudson, 2002) can only simulate short segments under the influence of recombination. The second challenge facing simulation methods is that sample sizes in genetic studies have grown very quickly in recent years, enabled by the precipitous fall in genome sequencing costs. Human datasets like the UK Biobank (Bycroft et al., 2018) and gnomAD (Karczewski et al., 2020) now consist of hundreds of thousands of genomes and many other datasets on a similar scale are becoming available (Tanjo et al., 2021). Classical simulators such as `ms` and even fast approximate methods such as `scrm` (Staab et al., 2015) simply cannot cope with such a large number of samples.

The `msprime` simulator (Kelleher et al., 2016; Kelleher and Lohse, 2020) has greatly increased the scope of coalescent simulations, and it is now straightforward to simulate millions of whole chromosomes for a wide range of organisms. The “succinct tree sequence” data structure (Kelleher et al., 2016, 2018, 2019; Wohns et al., 2021), originally introduced as part of `msprime`, makes it possible to store such large simulations in a few gigabytes, several orders of magnitude smaller than commonly used formats. The succinct tree sequence has also led to major advances in forwards-time simulation (Kelleher et al., 2018; Haller et al., 2018), ancestry inference (Kelleher et al., 2019; Wohns et al., 2021) and calculation of population genetic statistics (Kelleher et al., 2016; Ralph et al., 2020). Through a rigorous open-source community development process, `msprime` has gained a large number of features since its introduction, making it a highly efficient and flexible platform for population genetic simulation. This paper marks the release of `msprime` 1.0. We provide an overview of its extensive features, demonstrate its performance advantages over alternative software, and discuss opportunities for ongoing open-source community-based development.

The efficiency of coalescent simulations depends crucially on the assumption of neutrality, and it is important to note that there are many situations in which this will be a poor approximation of biological reality (Johri et al., 2021). In particular, background selection has been shown to affect genomewide sequence variation in a wide range of species (Charlesworth et al., 1993, 1995; Charlesworth and Jensen, 2021). Thus care must be taken to ensure that the results of purely neutral simulations are appropriate for the question and genomic partition under study. A major strength of `msprime`, however, is that it can be used in conjunction with forwards-time simulators, enabling the simulation of more realistic models than otherwise possible (Kelleher et al., 2018; Haller et al., 2018).

## Results

In the following sections we describe the main features of `msprime` 1.0, focusing on the aspects that are either new for this version, or in which our approach differs significantly from classical methods (summarised in Table 1). Where appropriate, we benchmark `msprime` against other simulators, but the comparisons are illustrative and not intended to be systematic or exhaustive. Please see Kelleher et al. (2016) for a performance comparison of `msprime` against simulators such as `ms, msms`, and `scrm`.

**Table 1:** Major features of `msprime` 1.0 added since version 0.3.0 (Kelleher et al., 2016).

|  |  |
| --- | --- |
| Interface | Separation of ancestry and mutation simulations. Ability to store arbitrary metadata along with simulation results, and automatic recording of provenance information for reproducibility. Jupyter notebook (Kluyver et al., 2016) integration. Rich suite of analytical and visualisation methods via `tskit` library. |
| Ancestry | SMC, SMC', Beta- and Dirac-coalescent, discrete time Wright-Fisher, and selective sweep models. Instantaneous bottlenecks. Discrete or continuous genomic coordinates, arbitrary ploidy, gene conversion. Output full ARG with recombination nodes, ARG likelihood calculations. Record full migration history and census events. Improved performance for large numbers of populations. Integration with forward simulators such as SLiM and `fwdpy11` ("recapitation"). |
| Demography | Improved interface with integrated metadata and referencing populations by name. Import from Newick species tree, *BEAST (Heled and Drummond, 2009), and Demes (Gower et al., 2022). Numerical methods to compute coalescence rates. |
| Mutations | JC69, HKY, F84, GTR, BLOSUM62, PAM, infinite alleles, SLiM and general matrix mutation models. Varying rates along the genome, recurrent/back mutations, discrete or continuous genomic coordinates, overlaying multiple layers of mutations, exact times associated with mutations. |

### User interface

The majority of simulation packages are controlled either through a command line interface (e.g. Hudson, 2002; Kern and Schrider, 2016), a text-based input file format (e.g. Guillaume and Rougemont, 2006; Excoffier and Foll, 2011; Shlyakhter et al., 2014), or a mixture of both. Command line interfaces make it easy to run simple simulations, but as model complexity and the number of parameters increase, they become difficult to understand and error-prone (Ragsdale et al., 2020; Gower et al., 2022). Specifying parameters through a text file alleviates this problem to a degree, but lacks flexibility, for example, when running simulations with parameters drawn from a distribution. In practice, for any reproducible simulation project users will write a script to generate the required command lines or input parameter files, invoke the simulation engine, and process the results in some way. This process is cumbersome and labour intensive, and a number of packages have been developed to allow simulations to be run directly in a high-level scripting language (Staab and Metzler, 2016; Parobek et al., 2017; Gladstein et al., 2018).

The more recent trend has been to move away from this file and command-line driven approach and to instead provide direct interfaces to the simulation engines via an Application Programming Interface (API) (e.g. Thornton, 2014; Kelleher et al., 2016; Becheler et al., 2019; Haller and Messer, 2019). The primary interface for `msprime` is through a thoroughly documented Python API, which has encouraged the development of an ecosystem of downstream tools (Terhorst et al., 2017; Chan et al., 2018; Spence and Song, 2019; Adrion et al., 2020a,b; Kamm et al., 2020; McKenzie and Eaton, 2020; Montinaro et al., 2020; Terasaki Hart et al., 2021; Rivera-Colón et al., 2021). As well as providing a stable and efficient platform for building downstream applications, `msprime`’s Python API makes it much easier to build reproducible simulation pipelines, as the entire workflow can be encapsulated in a single script, and package and version dependencies explicitly stated using the `pip` or `conda` package managers. For example, the errors made in the influential simulation analysis of Martin et al. (2017) were only detected because the pipeline could be easily run and reanalysed (Ragsdale et al., 2020; Martin et al., 2020).

A major change for the `msprime` 1.0 release is the introduction of a new set of APIs, designed in part to avoid sources of error (see the Demography section) but also to provide more appropriate defaults while keeping compatibility with existing code. In the new APIs, ancestry and mutation simulation are fully separated (see Fig. 1), with the `sim_ancestry` and `sim_mutations` functions replacing the legacy simulate function. Among other changes, the new APIs default to discrete genome coordinates and finite sites mutations, making the default settings more realistic and resolving a major source of confusion and error. The previous APIs are fully supported and tested, and will be maintained for the foreseeable future. The `msp` program (a command line interface to the library) has been extended to include new commands for simulating ancestry and mutations separately. A particularly useful feature is the ability to specify demographic models in Demes format (Gower et al., 2022) from the command line, making simulation of complex demographies straightforward. We also provide an `ms` compatible command line interface to support existing workflows.

**Figure 1:**
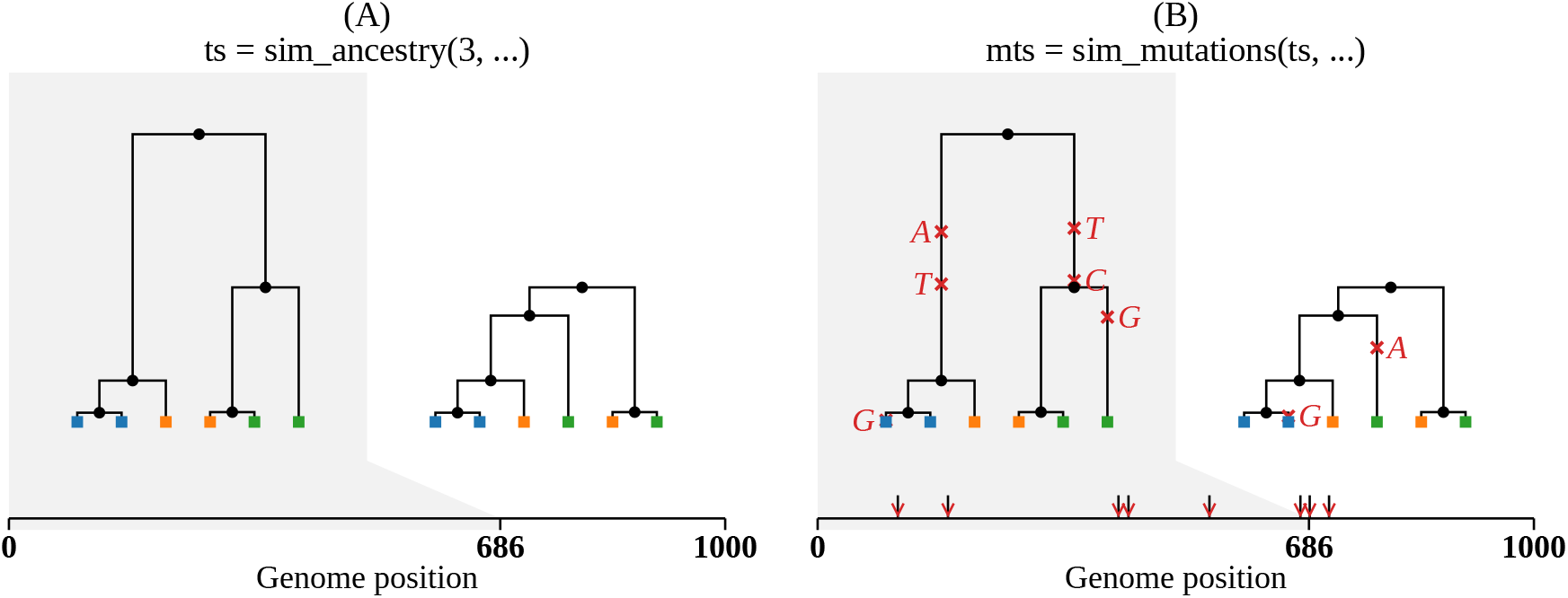
Visualisation of the separation between ancestry and mutation simulation. (A) The result of an invocation of `sim_ancestry` is two trees along a 1kb chunk of genome relating three diploid samples. Each diploid individual consists of two genomes (or nodes), indicated by colour. (B) This ancestry is provided as the input to `sim_mutations`, which adds mutations. Graphics produced using `tskit`’s `draw_svg` method.

### Tree sequences

One of the key reasons for `msprime`’s substantial performance advantage over other simulators (Kelleher et al., 2016) is its use of the “succinct tree sequence” data structure to represent simulation results. The succinct tree sequence (usually abbreviated to “tree sequence”) was introduced by Kelleher et al. (2016) to concisely encode genetic ancestry and sequence variation and was originally implemented as part of `msprime`. We subsequently extracted the core tree sequence functionality from `msprime` to create the `tskit` library, which provides a large suite of tools for processing genetic ancestry and variation data via APIs in the Python and C languages (Tskit developers, 2022). The availability of `tskit` as a liberally licensed (MIT) open source toolkit has enabled several other projects (e.g. Kelleher et al., 2019; Haller and Messer, 2019; Wohns et al., 2021; Terasaki Hart et al., 2021) to take advantage of the same efficient data structures used in `msprime`, and we hope that many more will follow. While a full discussion of tree sequences and the capabilities of `tskit` is beyond the scope of this article, we summarise some aspects that are important for simulation.

Let us define a genome as the complete set of genetic material that a child inherits from one parent. Thus, a diploid individual has two (monoploid) genomes, one inherited from each parent. Since each diploid individual lies at the end of two distinct lineages of descent, they will be represented by *two* places (nodes) in any genealogical tree. In the tree sequence encoding a *node* therefore corresponds to a single genome, which is associated with its creation time (and other op-tional information), and recorded in a simple tabular format (Fig. 2). Genetic inheritance between genomes (nodes) is defined by edges. An *edge* consists of a parent node, a child node and the left and right coordinates of the contiguous chromosomal segment over which the child genome inherited genetic material from the parent genome. Parent and child nodes may correspond to ancestor and descendant genomes separated by many generations. Critically, edges can span multiple trees along the genome (usually referred to as “marginal” trees), and identical node IDs across different trees corresponds to the same ancestral genome. For example, in Fig. 2 the branch from node 0 to 4 is present in both marginal trees, and represented by a single edge (the first row in the edge table). This simple device, of explicitly associating tree nodes with specific ancestral genomes and recording the contiguous segments over which parent-child relationships exist, generalises the original “coalescence records” concept (Kelleher et al., 2016), and is the key to the efficiency of tree sequences (Kelleher et al., 2018, 2019; Ralph et al., 2020). Note that this formulation is fully compatible with the concept of an Ancestral Recombination Graph (ARG) and any ARG topology can be fully and efficiently encoded in the node and edge tables illustrated in Fig. 2; see the Ancestral Recombination Graphs section below for more details.

**Figure 2:**
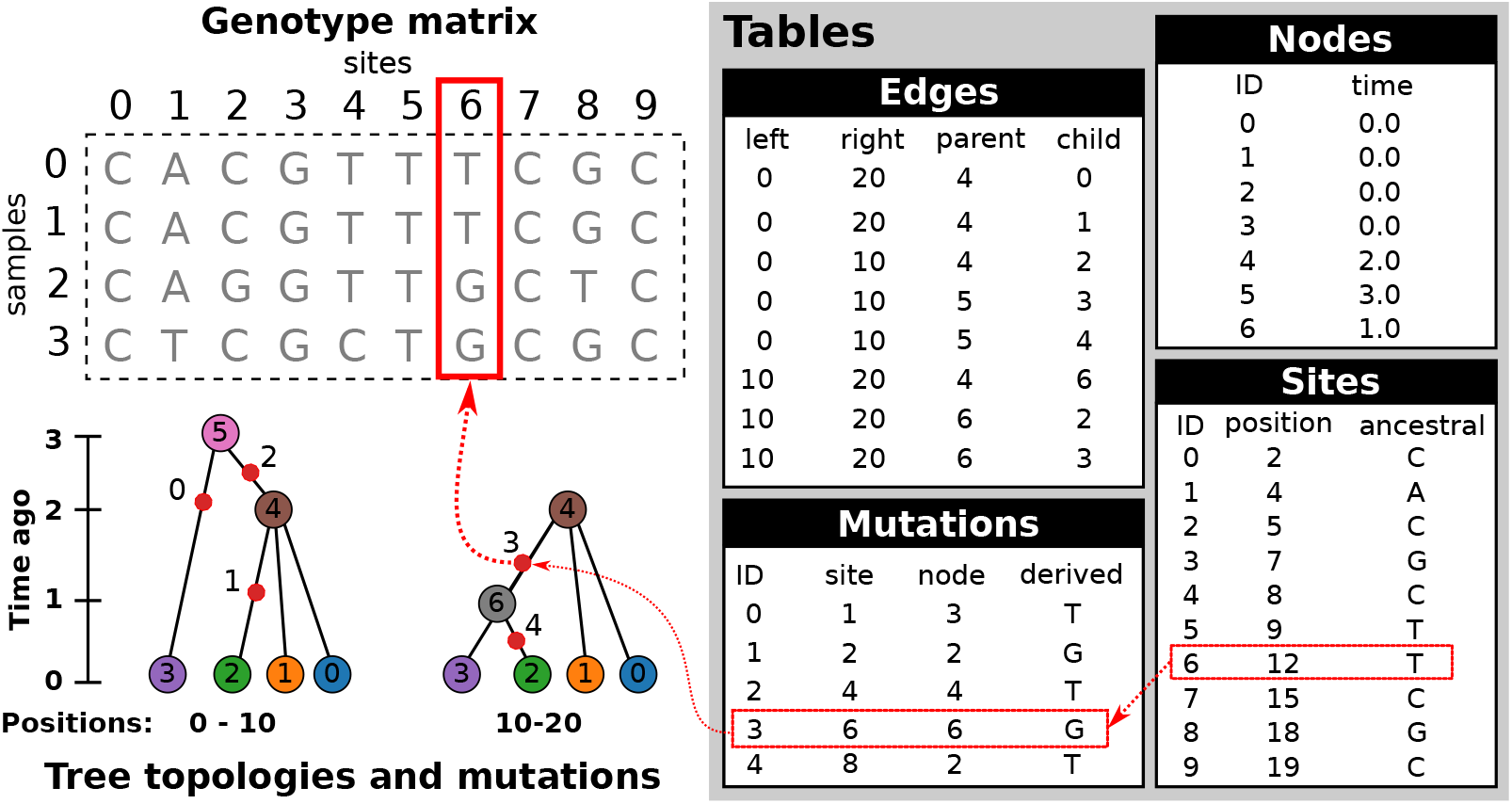
An example tree sequence describing genealogies and sequence variation for four samples at ten sites on a chromosome of twenty bases long. Information is stored in a set of tables (the tables shown here include only essential columns, and much more information can be associated with the various entities). The node table stores information about sampled and ancestral genomes. The edge table describes how these genomes are related along a chromosome, and defines the genealogical tree at each position. The site and mutation tables together describe sequence variation among the samples. The genotype matrix and tree topologies shown on the left are derived from these tables.

The final output of most population genetic simulations is some representation of sequence variation among the specified samples. For coalescent simulations, we usually have three steps: (1) simulate the genetic ancestry, and optionally output the resulting marginal trees; (2) simulate sequence evolution conditioned on this ancestry by generating mutations (see the Simulating mutations section); and (3) output the resulting nucleotide sequences by percolating the effects of the mutations through the trees. Information about the mutations themselves—e.g., where they have occurred on the trees—is usually not retained or made available for subsequent analysis. In `msprime`, however, we skip step (3), instead using `tskit`’s combined data model of ancestry and mutations to represent the simulated sequences. As illustrated in Fig. 2, mutations are a fully integrated part of `tskit`’s tree sequence data model, and genetic variation is encoded by recording sites at which mutations have occurred, and where each mutation at those sites has occurred on the marginal tree. Crucially, the genome sequences themselves are never stored, or indeed directly represented in memory (although `tskit` can output the variant matrix in various formats, if required). It may at first seem inconvenient to have only this indirect representation of the genome sequences, but it is extremely powerful. Firstly, the storage space required for simulations is dramatically reduced. For a simulation of n samples with *m* variant sites, we would require *O*(*nm*) space to store the sequence data as a variant matrix. However, if this simulation was of a recombining genome with *t* trees, then the `tskit` tree sequence encoding requires *O*(*n* + *t* + *m*) space, assuming we have *O*(1) mutations at each site (Kelleher et al., 2016). For large sample sizes, this difference is profound, making it conceivable, for example, to store the genetic ancestry and variation data for the entire human population on a laptop (Kelleher et al., 2019). As well as the huge difference in storage efficiency, it is often far more efficient to compute statistics of the sequence data from the trees and mutations than it is to work with the sequences themselves. For example, computing Tajima’s *D* from simulated data stored in the `tskit` format is several orders of magnitude faster than efficient variant matrix libraries for large sample sizes (Ralph et al., 2020).

The vast genomic datasets produced during the SARS-CoV-2 pandemic have highlighted the advantages of storing genetic variation data using the underlying trees. Turakhia et al. (2021) propose the Mutation Annotated Tree (MAT) format (consisting of a Newick tree and associated mutations in a binary format) and the `matUtils` program as an efficient way to store and process large viral datasets (McBroome et al., 2021), achieving excellent compression and processing performance. Similarly, `phastsim` (De Maio et al., 2021) was developed to simulate sequence evolution on such large SARS-CoV-2 phylogenies, and also outputs a Newick tree annotated with mutations (not in MAT format) to avoid the bottleneck of generating and storing the simulated sequences. While these methods illustrate the advantages of the general approach of storing ancestry and mutations rather than sequences, they do not generalise beyond their immediate settings, and no software library support is available.

The software ecosystem built around `tskit` is stable, mature and rapidly growing. Simulators such as `fwdpy11` (Thornton, 2014), `SLiM` (Haller and Messer, 2019), `stdpopsim` (Adrion et al., 2020a), `Geonomics` (Terasaki Hart et al., 2021) and `GSpace` (Virgoulay et al., 2021), and inference methods such as `tsinfer` (Kelleher et al., 2019), `tsdate` (Wohns et al., 2021) and `Relate` (Speidel et al., 2019) use either the Python or C APIs to support outputting results in tree sequence format. Tree sequences are stored in an efficient binary file format, and are fully portable across operating systems and processor architectures. The `tskit` library ensures interoperability between programs by having strict definitions of how the information in each of the tables is interpreted, and stringent checks for the internal consistency of the data model.

### Data analysis

The standard way of representing simulation data is to render the results in a text format, which must subsequently be parsed and processed as part of some analysis pipeline. For example, `ms` outputs a set of sequences and can also optionally output the marginal trees along the genome in Newick format, and variants of this approach are used by many simulators. Text files have many advantages, but are slow to process at scale. The ability to efficiently process simulation results is particularly important in simulation-based inference methods such as Approximate Bayesian Computation (ABC) (Beaumont et al., 2002; Csilléry et al., 2010; Wegmann et al., 2010) and machine learning based approaches (Sheehan and Song, 2016; Chan et al., 2018; Schrider and Kern, 2018; Flagel et al., 2019; Sanchez et al., 2020). Clearly, simulation efficiency is crucial since the size and number of simulations that can be performed determines the depth to which one can sample from the model and parameter space. Equally important, however, is the efficiency with which the simulation results can be transformed into the specific input required by the inference method. In the case of ABC, this is usually a set of summary statistics of the sequence data, and methods avoid the bottleneck of parsing text-based file formats to compute these statistics by either developing their own simulators (e.g. Cornuet et al., 2008; Lopes et al., 2009) or creating forked versions (i.e., modified copies) of existing simulators (e.g. Thornton and Andolfatto, 2006; Hickerson et al., 2007; Pavlidis et al., 2010; Huang et al., 2011; Quinto-Cortés et al., 2018), tightly integrated with the inference method. Modern approaches to ABC such as ABC-RF (Raynal et al., 2019; Pudlo et al., 2016) and ABC-NN (Csilléry et al., 2012; Blum and François, 2010) use large numbers of weakly informative statistics, making the need to efficiently compute statistics from simulation results all the more acute. By using the stable APIs and efficient data interchange mechanisms provided by `tskit`, the results of an `msprime` simulation can be immediately processed, without format conversion overhead. The `tskit` library has a rich suite of population genetic statistics and other utilities, and is in many cases orders of magnitude faster than matrix-based methods for large sample sizes (Ralph et al., 2020). Thus, the combination of `msprime` and `tskit` substantially increases the overall efficiency of many simulation analysis pipelines.

Classical text based output formats like `ms` are inefficient to process, but also lack a great deal of important information about the simulated process. The tree-by-tree topology information output by simulators in Newick format lacks any concept of node identity, and means that we cannot reliably infer information about ancestors from the output. Because Newick stores branch lengths rather than node times, numerical precision issues also arise for large trees (McGill et al., 2013). Numerous forks of simulators have been created to access information not provided in the output. For example, `ms` has been forked to output information about migrating segments (Rosenzweig et al., 2016), ancestral lineages (Chen and Chen, 2013), and `ms`’s fork `msHOT` (Hellenthal and Stephens, 2007) has in turn been forked to output information on local ancestry (Racimo et al., 2017). All of this information is either directly available by default in `msprime`, or can be optionally stored via options such as `record_migrations` or `record_full_arg` (see the Ancestral Recombination Graphs section) and can be efficiently and conveniently processed via `tskit` APIs.

### Simulating mutations

Because coalescent simulations are usually concerned with neutral evolution (see the Selective sweeps section, however) the problem of generating synthetic genetic variation can be decomposed into two independent steps: firstly, simulating genetic ancestry (the trees), then subsequently simulating variation by superimposing mutation processes on those trees (see Fig. 1). A number of programs exist to place mutations on trees: for instance, the classical `Seq-Gen` program (Rambaut and Grassly, 1997) supports a range of different models of sequence evolution, and various extensions to the basic models have been proposed (e.g. Cartwright, 2005; Fletcher and Yang, 2009). Partly for efficiency and partly in the interest of simplicity for users (i.e., to avoid intermediate text format conversions), population genetic simulators have tended to include their own implementations of mutation simulation, with most supporting the infinite sites model (e.g. Hudson, 2002) but with several supporting a wide range of different models of sequence evolution (e.g. Mailund et al., 2005; Excoffier and Foll, 2011; Virgoulay et al., 2021). Thus, despite the logical separation between the tasks of simulating ancestry and neutral sequence evolution, the two have been conflated in practice.

Part of the reason for this poor record of software reuse and modularity is the lack of standardised file formats, and in particular, the absence of common library infrastructure to abstract the details of interchanging simulation data. Although `msprime` also supports simulating both ancestry and mutations, the two aspects are functionally independent within the software; both ancestry and mutation simulators are present in `msprime` for reasons of convenience and history, and could be split into separate packages. The efficient C and Python interfaces for `tskit` make it straight-forward to add further information to an existing file, and because of its efficient data interchange mechanisms, there is no performance penalty for operations being performed in a different software package. Thanks to this interoperability, `msprime`’s mutation generator can work with *any* `tskit` tree sequence, be it simulated using `SLiM` (Haller and Messer, 2019) or `fwdpy11` (Thornton, 2014), or estimated from real data (Kelleher et al., 2019; Speidel et al., 2019; Wohns et al., 2021). It is a modular component intended to fit into a larger software ecosystem, and is in no way dependent on `msprime`’s ancestry simulator.

We have greatly extended the sophistication of `msprime`’s mutation generation engine for version 1.0, achieving near feature-parity with `Seq-Gen`. We support a large number of mutation models, including the JC69 (Jukes et al., 1969), F84 (Felsenstein and Churchill, 1996), and GTR (Tavaré et al., 1986) nucleotide models and the BLOSUM62 (Henikoff and Henikoff, 1992) and PAM (Dayhoff et al., 1978) amino acid models. Other models, such as the Kimura two and three parameter models (Kimura, 1980, 1981), can be defined easily and efficiently in user code by specifying a transition matrix between any number of alleles. Mutation rates can vary along the genome, and multiple mutation models can be imposed on a tree sequence by overlaying mutations in multiple passes. We have extensively validated the results of mutation simulations against both theoretical expectations and output from `Seq-Gen` (Rambaut and Grassly, 1997) and `Pyvolve` (Spielman and Wilke, 2015).

Simulating mutations in `msprime` is efficient. Fig. 3 shows the time required to generate mutations (using the default JC69 model) on simulated tree sequences for a variety of mutation rates as we vary the number of samples (Fig. 3A) and the sequence length (Fig. 3B). For example, the longest running simulation in Fig. 3B required less than 2 seconds to generate an average of 1.5 million mutations over 137,081 trees in a tree sequence with 508,125 edges. This efficiency for large numbers of trees is possible because the tree sequence encoding allows us to generate mutations on an edge-by-edge basis (see Fig. 2 and the Mutation generation appendix), rather than tree-by-tree and branch-by-branch as would otherwise be required. Simulating mutations on a single tree is also very efficient; for example, we simulated mutations under the BLOSUM62 amino acid model for a tree with 10^6^ leaves over 10^4^ sites (resulting in ~260,000 mutations) in about 0.8 seconds, including the time required for file input and output. We do not attempt a systematic benchmarking of `msprime`’s mutation generation code against other methods, because at this scale it is difficult to disentangle the effects of inefficient input and output formats from the mutation generation algorithms. Given the above timings, it seems unlikely that generating mutations with `msprime` would be a bottleneck in any realistic analysis.

**Figure 3:**
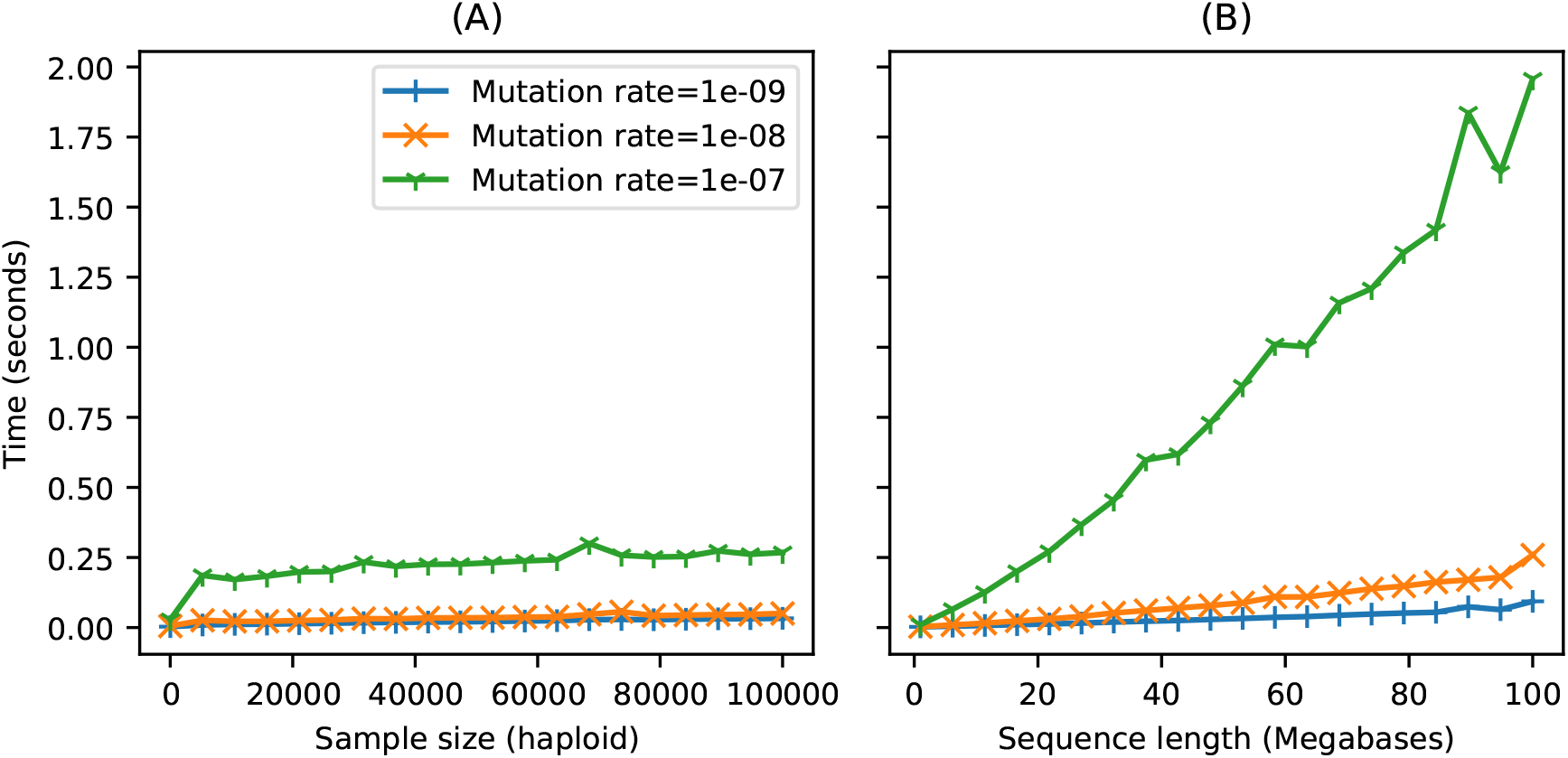
Time required to run `sim_mutations` on tree sequences generated by `sim_ancestry` (with a population size of 10^4^ and recombination rate of 10^−8^) for varying (haploid) sample size and sequence length. We ran 10 replicate mutation simulations each for three different mutation rates, and report the average CPU time required (Intel Core i7-9700). (A) Holding sequence length fixed at 10 megabases and varying the number of samples (tree tips) from 10 to 100,000. (B) Holding number of samples fixed at 1000, and varying the sequence length from 1 to 100 megabases.

There are many ways in which the mutation generation code in `msprime` could be extended. For example, we intend to add support for microsatellites (Mailund et al., 2005), codon models (Arenas and Posada, 2007) and indels (Cartwright, 2005; Fletcher and Yang, 2009), although changes may be required to `tskit`’s data model which is currently based on the assumption of independent sites.

### Recombination

Crossover recombination is implemented in `msprime` using Hudson’s algorithm, which works backwards in time, generating common ancestor and recombination events and tracking their effects on segments of ancestral material inherited from the sample (Hudson, 1983a, 1990; Kelleher et al., 2016). Common ancestor events merge the ancestral material of two lineages, and result in coalescences in the marginal trees when ancestral segments overlap. Recombination events split the ancestral material for some lineage at a breakpoint, creating two independent lineages. Using the appropriate data structures (Kelleher et al., 2016), this process is much more efficient to simulate than the equivalent left-to-right approach (Wiuf and Hein, 1999b,a). In `msprime` 1.0, recombination rates can vary along a chromosome, allowing us to simulate recombination hotspots and patterns of recombination from empirical maps. The implementation of recombination in `msprime` is extensively validated against analytical results (Hudson, 1983a; Kaplan and Hudson, 1985) and simulations by `ms`, `msHOT` and `SLiM`.

The Sequentially Markovian Coalescent (SMC) is an approximation of the coalescent with recombination (McVean and Cardin, 2005; Marjoram and Wall, 2006), and was primarily motivated by the need to simulate longer genomes than was possible using tools like `ms`. The SMC is a good approximation to the coalescent with recombination when we have fewer than five sampled genomes (Hobolth and Jensen, 2014; Wilton et al., 2015), but the effects of the approximation are less well understood for larger sample sizes, and several approaches have been proposed that allow simulations to more closely approximate the coalescent with recombination (Chen et al., 2009; Wang et al., 2014; Staab et al., 2015). The SMC and SMC’ models are supported in `msprime` 1.0. However, they are currently implemented using a naive rejection sampling approach, and are somewhat slower to simulate than the exact coalescent with recombination. These models are therefore currently only appropriate for studying the SMC approximations themselves, although we intend to implement them more efficiently in future versions.

In human-like parameter regimes and for large sample sizes, `msprime`’s implementation of the exact coalescent with recombination comprehensively outperforms all other simulators, including those based on SMC approximations (Kelleher et al., 2016). However, it is important to note that although the implementation of Hudson’s algorithm is very efficient, it is still quadratic in the population scaled recombination rate *ρ* = 4*N_e_L*, where *L* is the length of the genome in units of recombination distance. This is because Hudson’s algorithm tracks recombinations not only in segments ancestral to the sample, but also between ancestral segments. As mentioned above, common ancestor events in which the ancestral material of two lineages is merged only result in coalescences in the marginal trees if their ancestral segments overlap. If there is no overlap, the merged segments represent an ancestral chromosome that is a genetic ancestor of the two lineages, but not the most recent common genetic ancestor at any location along the genome. When this happens, the merged lineage carries “trapped” genetic material that is not ancestral to any samples, but where recombinations can still occur (Wiuf and Hein, 1999b). For large *ρ*, recombination events in trapped ancestral material will dominate, and so we can use this as a proxy for the overall number of events in Hudson’s algorithm. Hein et al. (2004, Eq. 5.10) gave

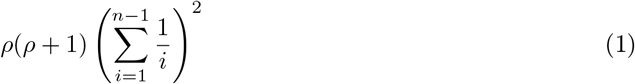

as an upper bound on the number of recombination events within trapped ancestral material for *n* samples. As discussed in the Time complexity of Hudson’s algorithm appendix, the quadratic dependence of simulation running time on *ρ* implied by (1) is well supported by observations, and provides a useful means of predicting how long a particular simulation might require.

### Gene conversion

Gene conversion is a form of recombination that results in the transfer of a short segment of genetic material, for example between homologous chromosomes (Chen et al., 2007). Since gene conversion impacts much shorter segments than crossover recombination (typically below 1kb) it affects patterns of linkage disequilibrium differently (Korunes and Noor, 2017). Wiuf and Hein (2000) modelled gene conversion in the coalescent via a rate at which gene conversion events are initiated along the genome and a geometrically distributed tract length. In terms of the ancestral process, gene conversion differs from crossover recombination (as described in the previous section) in that it extracts a short tract of ancestry into an independent lineage, rather than splitting ancestry to the left and right of a given breakpoint. We have implemented this model of gene conversion in `msprime` 1.0, and validated the output against `ms` and analytical results (Wiuf and Hein, 2000).

Gene conversion is particularly useful to model homologous recombination in bacterial evolution, and so we compare the performance of `msprime` with gene conversion to two specialised bacterial simulators, `SimBac` (Brown et al., 2016) and `fastSimBac` (De Maio and Wilson, 2017). Figure 4A shows that `msprime` is far more efficient than both `SimBac` and the SMC-based approximation `fastSimBac`. Figure 4B shows that `msprime` requires somewhat more memory than `fastSimBac`, (as expected since `fastSimBac` uses a left-to-right SMC approximation) but is still reasonably modest at around 1GiB for a simulation of 500 whole *E. coli* genomes. However, `msprime` is currently lacking many of the specialised features required to model bacteria, and so an important avenue for future work is to add features such as circular genomes and bacterial gene transfer (Baumdicker and Pfaffelhuber, 2014).

**Figure 4:**
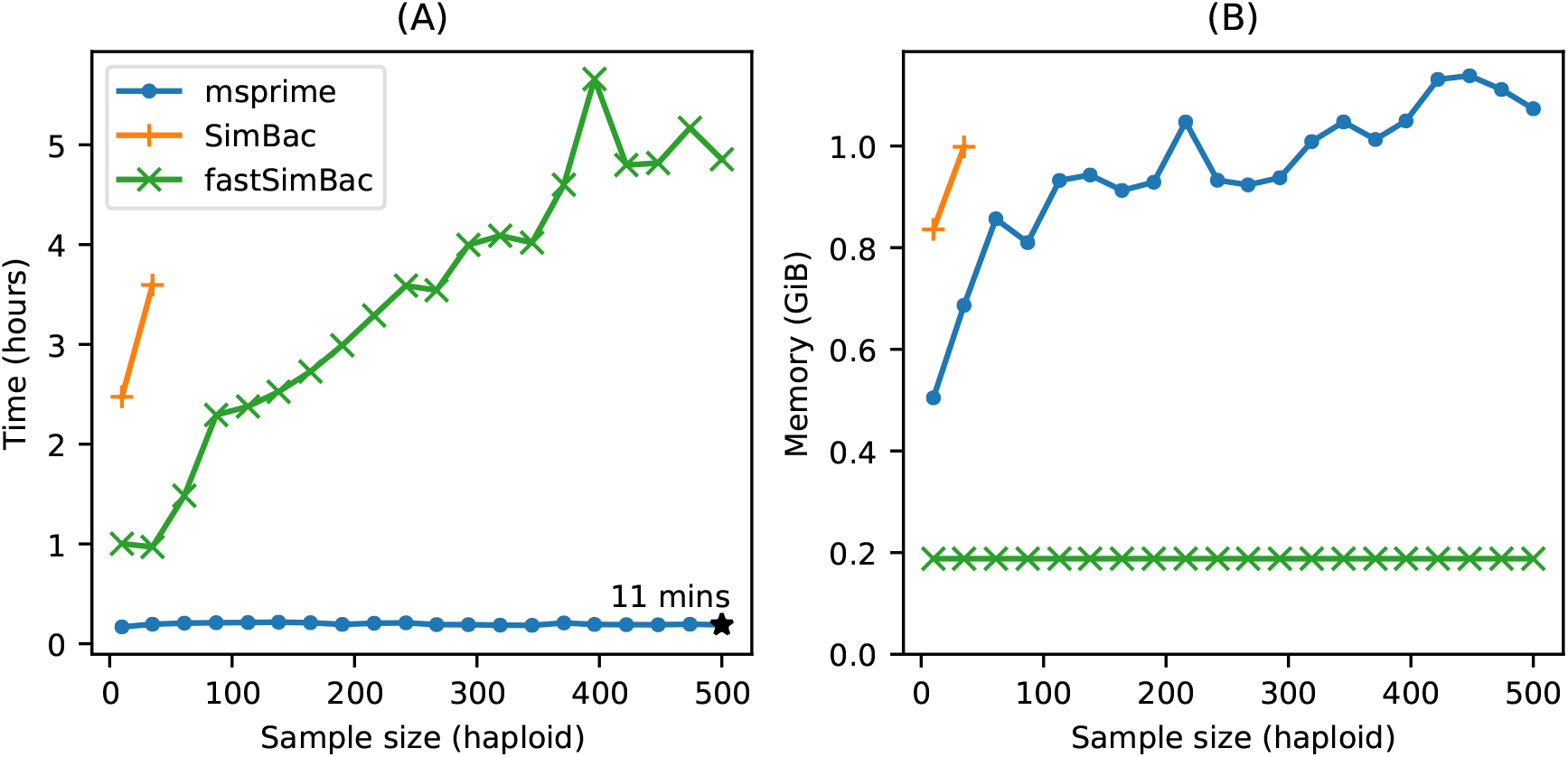
Comparison of simulation performance using `msprime` (`sim_ancestry`), SimBac, and `fastSimBac` for varying (haploid) sample sizes, and the current estimates for *E. coli* parameters (Lapierre et al., 2016): a 4.6Mb genome, *N_e_* = 1.8 × 10^8^, gene conversion rate of 8.9 × 10^-11^ per base and mean tract length of 542. We report (A) the total CPU time and (B) maximum memory usage averaged over 5 replicates (Intel Xeon E5-2680 CPU). We did not run `SimBac` beyond first two data points because of the very long running times.

### Demography

One of the key applications of population genetic simulations is to generate data for complex demographies. Beyond idealised cases such as stepping-stone or island models, or specialised cases such as isolation-with-migration models, analytical results are rarely possible. Simulation is therefore integral to the development and evaluation of methods for demographic inference. The demography model in `msprime` is directly derived from the approach used in `ms`, and supports an arbitrary number of randomly mating populations exchanging migrants at specified rates. A range of demographic events are supported, which allow for varying population sizes and growth rates, changing migration rates over time, as well as population splits, admixtures and pulse migrations.

A major change for `msprime` 1.0 is the introduction of the new Demography API, designed to address a design flaw in the `msprime` 0.x interface which led to avoidable errors in downstream simulations (Ragsdale et al., 2020). The new API is more user-friendly, providing the ability, for example, to refer to populations by name rather than their integer identifiers. We also provide numerical methods to compute the coalescence rates for two or more lineages which can be inverted to obtain the “inverse instantaneous coalescence rate” of Chikhi et al. (2018). Many popular approaches in population genetics use the distribution of coalescence rates between pairs of lineages to infer effective population sizes over time (Li and Durbin, 2011; Sheehan et al., 2013; Schiffels and Durbin, 2014) or split times and subsequent migration rates between populations (Wang et al., 2020). These numerical methods provide a valuable ground-truth when evaluating such inference methods, as illustrated by Adrion et al. (2020a).

### Instantaneous bottlenecks

A common approach to modelling the effect of demographic history on genealogies is to assume that effective population size (*N_e_*) changes in discrete steps which define a series of epochs (Griffiths et al., 1994; Marth et al., 2004; Keightley and Eyre-Walker, 2007; Li and Durbin, 2011). In this setting of piecewise constant *N_e_*, capturing a population bottleneck requires three epochs: *N_e_* is reduced by some fraction *b* at the start of the bottleneck, *T_start_*, and recovers to its initial value at time *T_end_* (Marth et al., 2004). If bottlenecks are short both on the timescale of coalescence and mutations, there may be little information about the duration of a bottleneck (*T_end_* – *T_start_*) in sequence data. Thus a simpler, alternative model is to assume that bottlenecks are instantaneous (*T_end_* – *T_start_* → 0) and generate a sudden burst of coalescence events (a multiple merger event) in the genealogy. The strength of the bottleneck *B* can be thought of as an (imaginary) time period during which coalescence events are collapsed, i.e. there is no growth in genealogical branches during *B* and the probability that a single pair of lineages entering the bottleneck coalesce during the bottleneck is 1 – *e^−B^*. Although this simple two parameter model of bottlenecks is attractive and both analytic results and empirical inference (Griffiths et al., 1994; Birkner et al., 2009; Galtier et al., 2000; Bunnefeld et al., 2015) have been developed under this model, there has been no software available to simulate data under instantaneous bottleneck histories.

We have implemented instantaneous bottlenecks in `msprime` 1.0 using a variant of Hudson’s linear time single-locus coalescent algorithm (Hudson, 1990), and validated the results by comparing against analytical expectations (Bunnefeld et al., 2015).

### Multiple merger coalescents

Kingman’s coalescent assumes that only two ancestral lineages can merge at each merger event. Although this is generally a reasonable approximation, there are certain situations in which the underlying mathematical assumptions are violated. For example in certain highly fecund organisms (Hedgecock, 1994; Beckenbach, 1994; Hedgecock and Pudovkin, 2011; Árnason, 2004; Irwin et al., 2016; Vendrami et al., 2021), where individuals have the ability to produce numbers of offspring on the order of the population size and therefore a few individuals may produce the bulk of the offspring in any given generation (Hedgecock, 1994). These population dynamics violate basic assumptions of the Kingman coalescent, and are better modelled by ‘multiple-merger’ coalescents (Donnelly and Kurtz, 1999; Pitman, 1999; Sagitov, 1999; Schweinsberg, 2000; Möhle and Sagitov, 2001), in which more than two lineages can merge in a given event. Multiple-merger coalescent processes have also been shown to be relevant for modelling the effects of selection on gene genealogies (Gillespie, 2000; Durrett and Schweinsberg, 2004; Desai et al., 2013; Neher and Hallatschek, 2013; Schweinsberg, 2017).

Although multiple merger coalescents have been of significant theoretical interest for around two decades, there has been little practical software available to simulate these models. Kelleher et al. (2013, 2014) developed packages to simulate a related spatial continuum model (Barton et al., 2010), Zhu et al. (2015) simulate genealogies within a species tree based on a multiple-merger model, and Becheler and Knowles (2020) provide a general method for simulating multiple merger processes as part of the Quetzal framework (Becheler et al., 2019). The `Beta-Xi-Sim` simulator (Koskela, 2018; Koskela and Wilke Berenguer, 2019) also includes a number of extensions to the Beta-coalescent. None of these methods work with large genomes, and very little work has been performed on simulating multiple merger processes with recombination.

We have added two multiple merger coalescent models in `msprime` 1.0, the Beta-coalescent (Schweinsberg, 2003) and “Dirac”-coalescent (Birkner et al., 2013a), allowing us to efficiently simulate such models with recombination for the first time. These simulation models have been extensively validated against analytical results from the site frequency spectrum (Birkner et al., 2013b; Blath et al., 2016; Hobolth et al., 2019) as well as more general properties of coalescent processes. See the Multiple merger coalescent model appendix for more details and model derivations.

### Ancestral Recombination Graphs

The Ancestral Recombination Graph (ARG) was introduced by Griffiths (Griffiths, 1991; Griffiths and Marjoram, 1997) to represent the stochastic process of the coalescent with recombination as a graph. This formulation is complementary to Hudson’s earlier work (Hudson, 1983a), and substantially increased our theoretical understanding of recombination. In Griffiths’ ARG formulation, a realisation of the coalescent with recombination is a graph in which vertices represent common ancestor or recombination events, and edges represent lineages. There is the “big” ARG, in which we track lineages arising out of recombinations regardless of whether they carry ancestral material (Ethier and Griffiths, 1990), and the “little” ARG in which we only track genetic ancestors. Over time, usage of the term has shifted away from its original definition as a stochastic process, to being interpreted as a representation of a particular genetic ancestry as a graph, without necessarily following the specific details of the Griffiths formulation (e.g. Minichiello and Durbin, 2006; Mathieson and Scally, 2020). Under the latter interpretation, the tree sequence encoding of genetic ancestry (described above) clearly *is* an ARG: the nodes and edges define a graph in which edges are annotated with the set of disjoint genomic intervals through which ancestry flows.

For our purposes, an ARG is a realisation of the coalescent with recombination, in the Griffiths (little ARG) sense. As described in detail by Kelleher et al. (2016), Hudson’s algorithm works by dynamically traversing a little ARG. The graph is not explicitly represented in memory, but is partially present through the extant lineages and the ancestral material they carry over time. We do not output the graph directly, but rather store the information required to recover the genealogical history as nodes and edges in a tree sequence. This is far more efficient than outputting the simulated ARG in its entirety. For a given scaled recombination rate *ρ* (setting aside the dependency on the sample size *n*) we know from Eq. (1) that the number of nodes in an ARG is *O*(*ρ*^2^), whereas the size of the tree sequence encoding is *O*(*ρ*) (Kelleher et al., 2016). This difference between a quadratic and a linear dependency on *ρ* is profound, and shows why large simulations cannot output an ARG in practice.

Although by default `msprime` outputs tree sequences that contain full information about the genealogical trees, their correlation structure along the chromosome, and the ancestral genomes on which coalescences occurred, some information is lost in this mapping down from ARG space to the minimal tree sequence form. In particular, we lose information about ancestral genomes that were common ancestors but in which no coalescences occurred, and also information about the precise time and chromosomal location of recombination events. In most cases, such information is of little relevance as it is in principle unknowable, but there are occasions such as visualisation or computing likelihoods (see below) in which it is useful. We therefore provide the `record_full_arg` option in `msprime` to store a representation of the complete ARG traversed during simulation. This is done by storing extra nodes (marked with specific flags, so they can be easily identified later) and edges in the tree sequence (Fig. 5). One situation in which a record of the full ARG is necessary is when we wish to compute likelihoods during inference. The likelihood is a central quantity in evaluating the plausibility of a putative ancestry as an explanation of DNA sequence data, both directly through e.g. approaches based on maximum likelihood, and as an ingredient of methods such as Metropolis-Hastings (Kuhner et al., 2000; Nielsen, 2000; Wang and Rannala, 2008). We provide functions to compute the likelihood of ARG realisations and mutational patterns under the standard coalescent and infinite sites mutation model. For details, see the appendix: Likelihood calculations.

**Figure 5:**
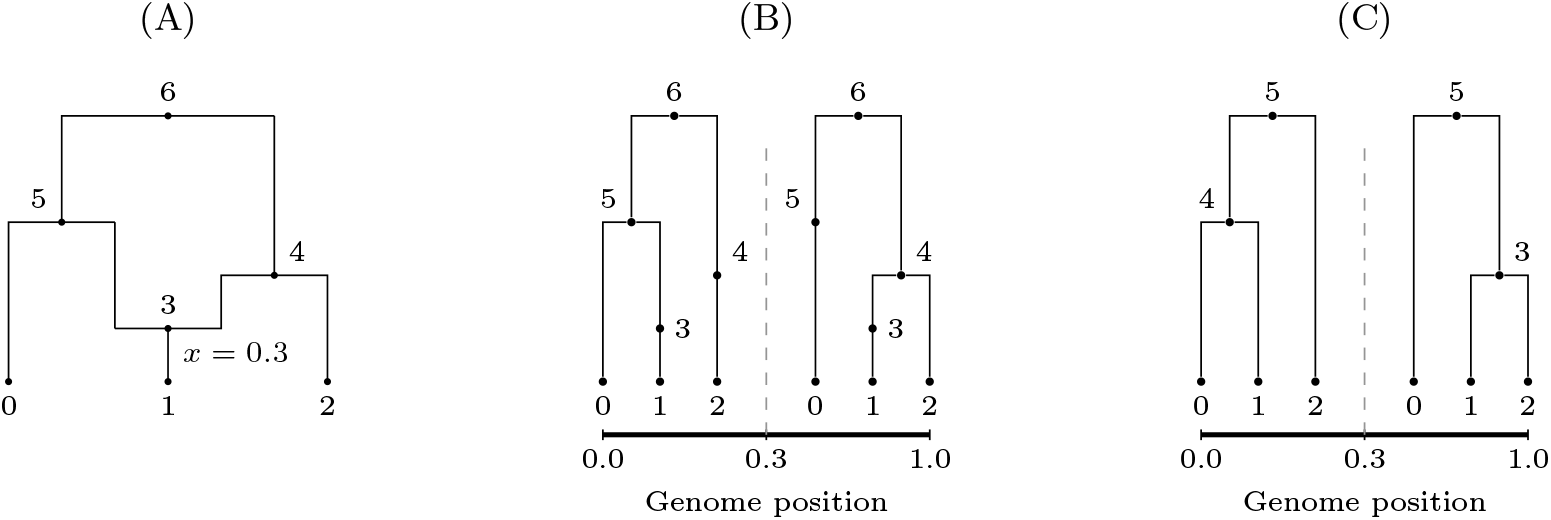
(A) A simple ARG in which a recombination occurs at position 0.3; (B) the equivalent topology depicted as a tree sequence, including the recombination node; (C) the same tree sequence topology “simplified” down to its minimal tree sequence representation. Note that the internal nodes have been renumbered in the simplified representation, so that, e.g., node 5 in (C) corresponds to node 6 in (A) and (B).

### Selective sweeps

Another elaboration of the standard neutral coalescent with recombination is the addition of selective sweeps (Kaplan et al., 1989; Braverman et al., 1995; Kim and Stephan, 2002). Sweeps are modelled by creating a structured population during the sojourn of the beneficial mutation through the population (i.e., the sweep phase) in which lineages may transit between favoured and unfavoured backgrounds through recombination. This approach allows for many selective sweep scenarios to be simulated efficiently, including recurrent, partial, and soft selective sweeps. However this efficiency comes at the cost of flexibility in comparison to forwards in time simulation. Several specialised simulators have been developed to simulate sweeps in the coalescent, including `SelSim` (Spencer and Coop, 2004), `mbs` (Teshima and Innan, 2009), `msms` (Ewing and Hermisson, 2010), `cosi2` (Shlyakhter et al., 2014) and `discoal` (Kern and Schrider, 2016).

Selective sweeps are implemented in the coalescent as a two step-process: first generating an allele frequency trajectory, and then simulating a structured coalescent process conditioned on that trajectory. Following `discoal`, we generate sweep trajectories in `msprime` using a jump process approximation to the conditional diffusion of an allele bound for fixation (Coop and Griffiths, 2004), as detailed in the Selective sweeps model appendix. Given a randomly generated allele frequency trajectory, the simulation of a sweep works by assigning lineages to two different structured coalescent “labels”, based on whether they carry the beneficial allele. The allele frequency trajectory determines the relative sizes of the “populations” in these labels over time, and therefore the rates at which various events occur. Common ancestor events can then only merge lineages from *within* a label, but lineages can transfer from one label to the other (i.e., from the advantageous to disadvantageous backgrounds, and vice versa) as a result of recombination events. Once we have reached the end of the simulated trajectory the sweep is complete, and we remove the structured coalescent labels. Simulation may then resume under any other ancestry model.

Fig. 6 compares the performance of `msprime` and `discoal` under a simple sweep model, and shows that `msprime` has far better CPU time and memory performance. Since our implementation uses the abstract label system mentioned above, adding support for similar situations, such as inversions (Peischl et al., 2013), should be straightforward.

**Figure 6:**
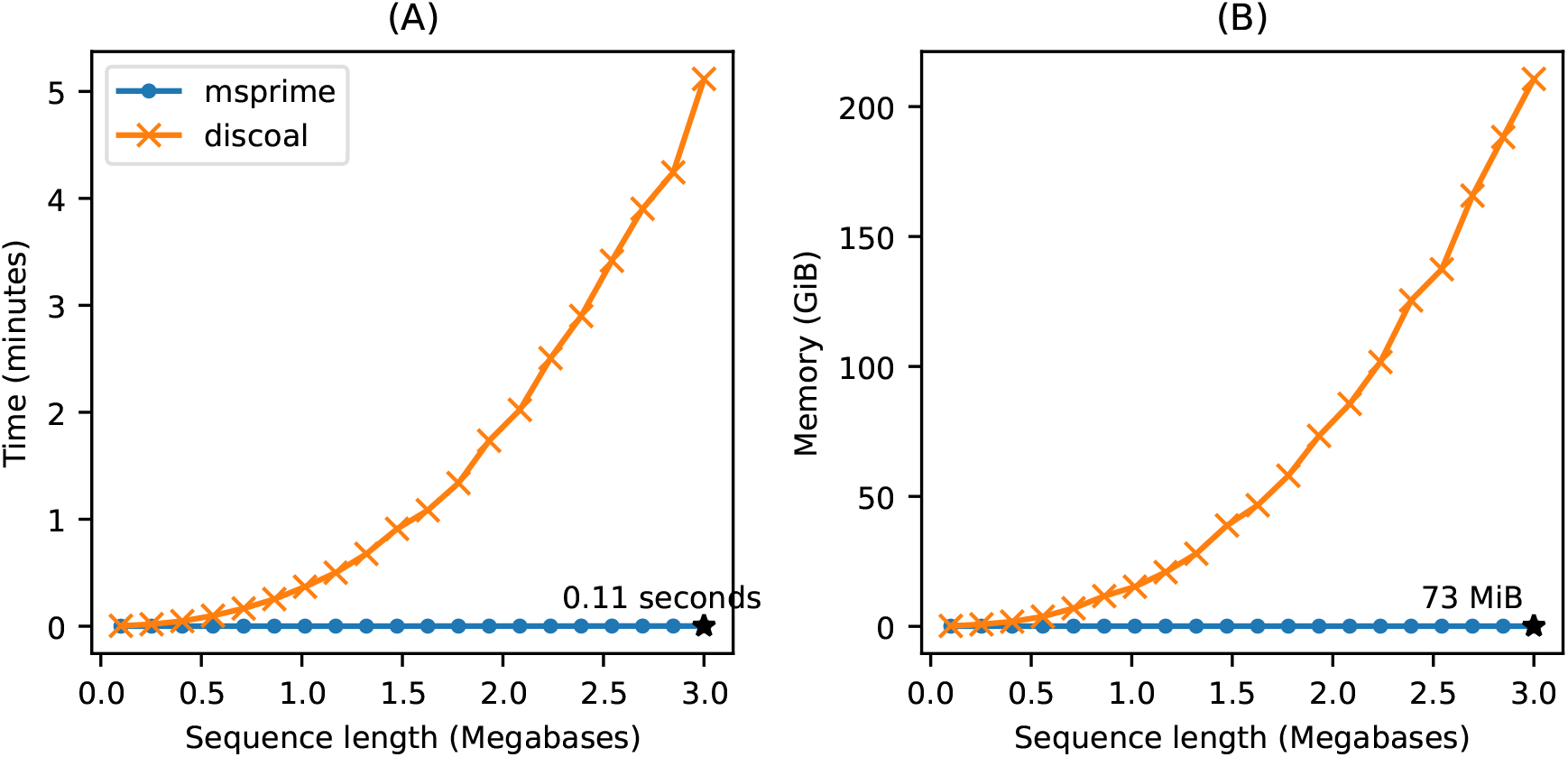
Comparison of selective sweep simulation performance in `msprime` (`sim_ancestry`) and discoal (Intel Xeon Gold 6148 CPU). We report the average CPU time and maximum memory usage when simulating 3 replicates for 100 diploid samples in a model with a single selective sweep in its history, where the beneficial allele had a selection coefficient of *s* = 0.05, a per-base recombination rate of 10^-8^, population size of *N* = 10^4^, and sequence length varying from 100kb–3000kb.

### Discrete time Wright-Fisher

The coalescent is an idealised model and makes many simplifying assumptions, but it is often surprisingly robust to violations of these assumptions (Wakeley et al., 2012). One situation in which the model does break down is the combination of large sample size and long recombining genomes, where the large number of recombination events in the recent past results in more than the biologically possible 2^*t*^ ancestors in *t* diploid generations (Nelson et al., 2020). This pathological behaviour results in identity-by-descent, long-range linkage disequilibrium and ancestry patterns deviating from Wright-Fisher expectations, and the bias grows with larger sample sizes (Wakeley et al., 2012; Bhaskar et al., 2014; Nelson et al., 2020). Precisely this problem occurs when simulating modern human datasets, and we have implemented a Discrete Time Wright-Fisher (DTWF) model in `msprime` to address the issue. The DTWF simulates backwards in time generation-by-generation so that each gamete has a unique diploid parent, and multiple recombinations within a generation results in crossover events between the same two parental haploid copies. The method is described in detail by Nelson et al. (2020).

Fig. 7 shows that `msprime` simulates the DTWF more quickly and requires substantially less memory than `ARGON` (Palamara, 2016), a specialised DTWF simulator. However, the generation-by-generation approach of the DTWF is less efficient than the coalescent with recombination when the number of lineages is significantly less than the population size (the regime where the coalescent is an accurate approximation), which usually happens in the quite recent past (Bhaskar et al., 2014). We therefore support changing the simulation model during a simulation so that we can run hybrid simulations, as proposed by Bhaskar et al. (2014). Any number of different simulation models can be combined, allowing for the flexible choice of simulation scenarios. As the DTWF improves accuracy of genealogical patterns in the recent past, we can simulate the recent history using this model and then switch to the standard coalescent to more efficiently simulate the more ancient history.

**Figure 7:**
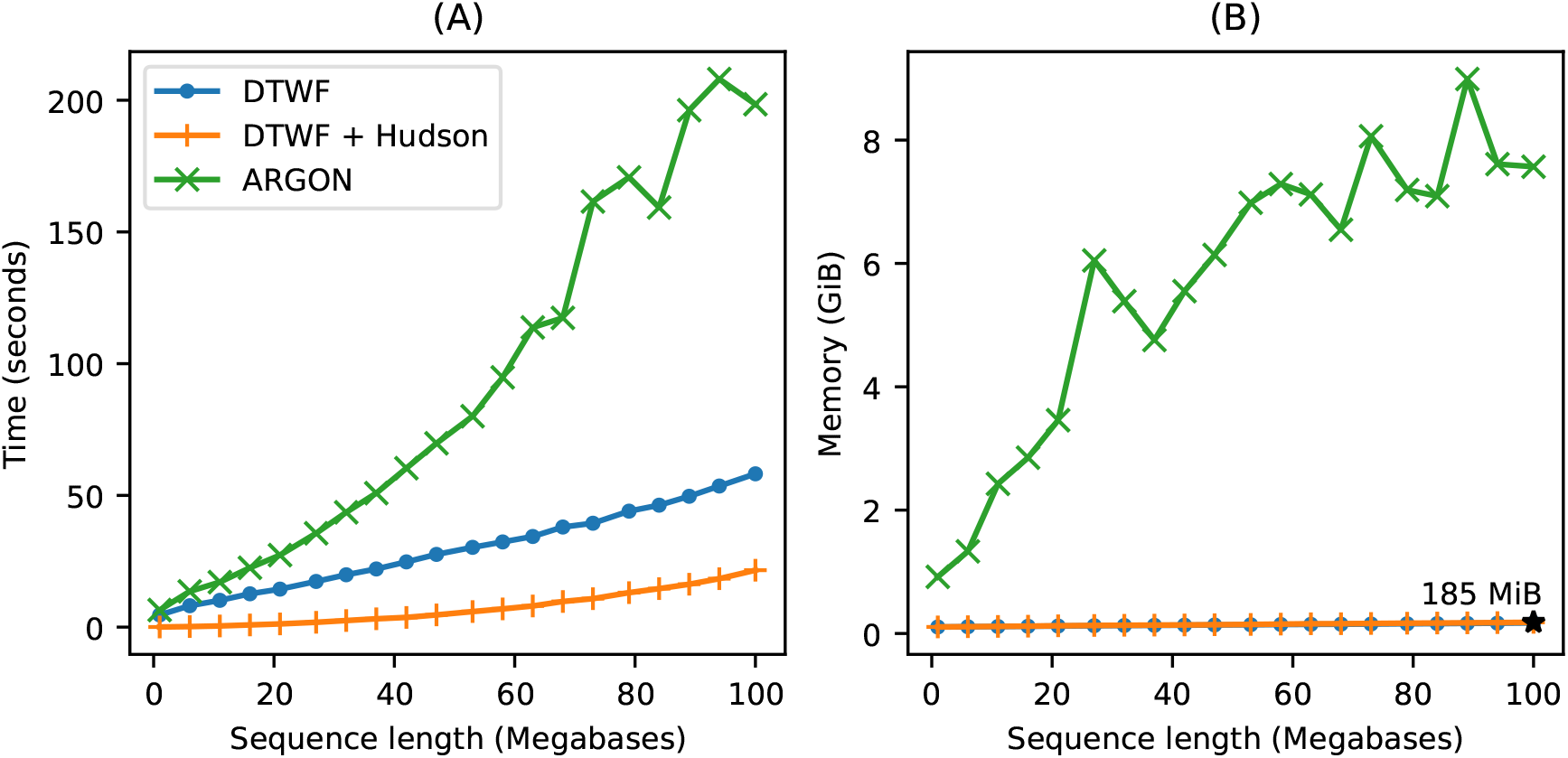
Comparison of Discrete Time Wright-Fisher (DTWF) simulation performance in `msprime` (`sim_ancestry`) and `ARGON` (Intel Xeon E5-2680 CPU). We ran simulations with a population size of 10^4^ and recombination rate of 10^−8^, with 500 diploid samples and varying sequence length. We report (A) total CPU time and (B) maximum memory usage; each point is the average over 5 replicate simulations. We show observations for `ARGON`, `msprime`’s DTWF implementation (“DTWF”) and a hybrid simulation of 100 generations of the DTWF followed by the standard coalescent with recombination (“DTWF + Hudson”). We ran `ARGON` with a mutation rate of 0 and with minimum output options, with a goal of measuring only ancestry simulation time. Memory usage for `msprime`’s DTWF and hybrid simulations are very similar.

### Integration with forward simulators

A unique feature of `msprime` is its ability to simulate genetic ancestries by extending an existing partial genetic ancestry. Given a tree sequence that is complete up until time *t* ago as input (where marginal trees may or may not have fully coalesced), `msprime` can efficiently obtain the segments of ancestral material present at this time, and then run the simulation backwards in time from there. This allows a simulated ancestry to be produced by any number of different processes across disjoint time slices. In practice this feature is used to “complete” forwards-time ancestry simulations (Kelleher et al., 2018) that may have not fully coalesced. This process (“recapitation”) can be orders of magnitude faster than the standard approach of neutral burn-in; see Haller et al. (2018) for more details and examples. This interoperability between simulators, where a partial ancestry simulation produced by `SLiM` (Haller and Messer, 2019) or `fwdpy11` (Thornton, 2014) can be picked up and completed by another simulator, with complete information retained—at scale—is unprecedented. There may be an opportunity for other forward genetic simulators (e.g. Gaynor et al., 2021) to leverage the tree sequence data format and associated tools.

### Development model

Msprime has a large number of features, encompassing the functionality of several more specialised simulators while maintaining excellent performance. It is developed by a geographically distributed team of volunteers under an open source community development model, with a strong emphasis on code quality, correctness, good documentation, and inclusive development. As in any large code base, unit tests play a key role in ensuring that new additions behave as expected and `msprime` has an extensive suite. These tests are run automatically on different operating systems on each pull request (where a contributor proposes a code change), using standard Continuous Integration (CI) methodology. Other CI services check for common errors, code formatting issues, and produce reports on the level of test coverage for the proposed change.

Unit tests are vital for ensuring software quality and correctness, but they are usually of little value in assessing the statistical properties of simulations. To validate the correctness of simulation output we maintain a suite of statistical tests (as of 1.0.0, 217 validation tests). These consist of running many replicate simulations to check the properties of the output against other simulators, and where possible against analytical results. For example, simulations of complex demography are validated against `ms`, selective sweeps against `discoal`, and Wright-Fisher simulations against forwards in time simulations in `SLiM`. This suite of tests is run before every release, to ensure that statistical errors have not been introduced.

More visibly to the end user, we also have a high standard for documentation, with precise, comprehensive, and cross-linked documentation that is automatically built from the code base and served through the website https://tskit.dev. With the goal of lowering the entry barrier to new users, we have invested significant effort in writing examples and introductions, and making common tasks discoverable. We also view contributions to documentation as equally important to the project as writing code or designing methods: what use would it be to write reliable, stable software if no-one used it?

An important goal of `msprime`’s development model is to maximise accessibility for prospective users and contributors, and to encourage diversity in our community. Gender and racial inequality caused by discrimination and marginalisation is a major problem across the sciences (Wellenreuther and Otto, 2016; Shannon et al., 2019) and in open source software development (Trinkenreich et al., 2021). Within our field, the contribution of women to early computational methods in population genetics was marginalised (Dung et al., 2019), and women continue to be under-represented in computational biology (Bonham and Stefan, 2017). The authorship of our paper reflects these trends, with a skew towards men and affiliations in the USA and Europe. We know the importance of creating and strengthening networks to develop and maintain a diverse community of contributors, and we are committed to fostering a supportive and collaborative environment that helps to address these inequalities in our field.

## Discussion

The 1.0 release of `msprime` marks a major increase in the breadth of available features and the potential biological realism of simulations. These abilities will allow researchers to perform more robust power analyses, more reliably test new methods, carry out more reliable inferences, and more thoroughly explore the properties of theoretical models. Despite this complexity and generality, `msprime`’s performance is state-of-the-art and all features are extensively tested and statistically validated. These advances have only been possible thanks to a distributed, collaborative model of software development, and the work of many people.

Even though simulation has long been a vital tool in population genetics, such collaborative software development has historically been uncommon. A huge proliferation of tools have been published (the references here are not exhaustive) and only a small minority of these are actively developed and maintained today. The ecosystem is highly fragmented, with numerous different ways of specifying parameters and representing results, and there are significant software quality issues at all stages. This is unsurprising, since the majority of simulation software development is performed by students, often without formal training in software development. The result resembles Haldane’s sieve for new mutations: many new pieces of software stay permanently on a dusty shelf of supplementary materials, while some of those that prove particularly useful when new (like dominant alleles) are quickly adopted. Although this has produced many good tools and enabled decades of research, it also represents a missed opportunity to invest as a community in shared infrastructure and mentorship in good software development practice.

Scientific software is vital apparatus, and must be engineered to a high quality if we are to trust its results. There is a growing realisation across the sciences (e.g. Siepel, 2019; Harris et al., 2020; Gardner et al., 2021) that investing in shared community infrastructure produces better results than a proliferation of individually maintained tools, allowing scientists to focus on their specific questions rather than software engineering. `Msprime` 1.0 is the result of such a community process, with features added by motivated users, taking advantage of the established development practices and infrastructure. Software development in a welcoming community, with mentorship by experienced developers, is a useful experience for many users. The skills that contributors learn can lead to greatly increased productivity in subsequent work (e.g., through more reliable code and better debugging skills). We hope that users who find that features they require are missing will continue to contribute to `msprime`, leading to a community project that is both high quality and sustainable in the long term.

The succinct tree sequence data structure developed for `msprime` provides a view of not only genetic variation, but also the genetic ancestry that produced that variation. Recent breakthroughs in methods to infer genetic ancestry in recombining organisms (Rasmussen et al., 2014; Kelleher et al., 2019; Speidel et al., 2019; Wohns et al., 2021; Schaefer et al., 2021; Speidel et al., 2021) have made it possible to estimate such ancestry from real data at scale for the first time (Harris, 2019; Tang, 2019). Given such inferred ancestry, many exciting applications become possible. For example, Osmond and Coop (2021) developed a method to estimate the location of genetic ancestors based on inferred trees, and other uses are sure to follow. Since the inferred genetic ancestry becomes the input for other downstream inferences, it is vitally important that these primary inferences are thoroughly validated, with the detailed properties of the inferred ancestries catalogued and understood. `Msprime` will continue to be an important tool for these inferences and validations, and in this context the ability to interoperate with other methods—particularly forwards simulators—through the succinct tree sequence data structure and `tskit` library will be essential.

## Availability

Msprime is freely available under the terms of the GNU General Public License v3.0, and can be installed from the Python Package Index (PyPI) or the conda-forge conda channel. Development is conducted openly on GitHub at https://github.com/tskit-dev/msprime/. The documentation for `msprime` is available at https://tskit.dev/msprime/docs/. The source code for all the evaluations and figures in this manuscript is available at https://github.com/tskit-dev/msprime-1.0-paper/.

## Acknowledgements

We acknowledge the contributions of Ivan Krukov who we consider eligible for authorship, but were unable to contact for approval. We would like to thank Iain Mathieson and Alywyn Scally for helpful comments on the manuscript. ADK was supported by NIH awards R01GM117241 and R01HG010774. BE was supported by DFG grant 273887127 through Priority Programme SPP 1819: Rapid Evolutionary Adaptation (grant STE 325/17-2) to Wolfgang Stephan; BE would also like to acknowledge funding through The Icelandic Research Centre (Rannís) through an Icelandic Research Fund Grant of Excellence nr. 185151-051 to Einar Árnason, Katrín Halldórsdóttir, Alison Etheridge, Wolfgang Stephan, and BE. FB is funded by the Deutsche Forschungsgemeinschaft EXC 2064/1 – Project number 390727645, and EXC 2124 – Project number 390838134. GB and KL are supported by an ERC starting grant (ModelGenomLand 757648) to KL. Graham Gower was supported by a Villum Fonden Young Investigator award to Fernando Racimo (project no. 00025300). Gregor Gorjanc is supported by the Chancellor’s Fellowship of the University of Edinburgh and the BBSRC grant to The Roslin Institute BBS/E/D/30002275. Jere Koskela is supported in part by EPSRC grant EP/R044732/1. Jerome Kelleher is supported by the Robertson Foundation. SG acknowledges funding from the Canada Research Chairs Program, from the Canadian Institutes of Health Research PJT 173300, and from the Canadian Foundation for Innovation.

## Appendix

### Mutation generation

The algorithm that `msprime` uses to simulate mutations on a tree sequence proceeds in two steps: first, mutations are “placed” on the tree sequence (i.e., sampling their locations in time, along the genome, and on the marginal tree), and then the ancestral and derived alleles of each mutation are generated. All mutation models share the code to place mutations, but choose alleles in different ways.

First, mutations are placed on the tree sequence under an inhomogeneous Poisson model by applying them independently to each edge. If an edge spans a region [*a, b*) of the genome and connected parent and child nodes with times *s* < *t*, and the mutation rate locally is *μ*, then the number of mutations on the edge is Poisson with mean *μ*(*t* – *s*)(*b* – *a*), and each mutation is placed independently at a position chosen uniformly in [*a, b*) and a time uniformly in [*s, t*). In a discrete genome, all positions are integers and so more than one mutation may occur at the same position on the same edge. Otherwise (i.e., for an infinite-sites model), positions are rejection sampled to obtain a unique floating-point number. If an edge spans a region of the genome with more than one mutation rate, this is done separately for each sub-region on which the mutation rate is constant. Since each edge is processed independently, the algorithm scales linearly with the number of edges in the tree sequence.

Next, alleles are chosen for each mutation. If the site was not previously mutated, then a new ancestral allele is chosen for the site, according to an input distribution of ancestral state allele probabilities. Then, each mutation on the tree is considered in turn, and a derived allele is randomly chosen based on the parental allele (which may be the ancestral allele or the derived allele of a previous mutation). Finally, information about the mutations are recorded in the site and mutation tables of the tree sequence.

A mutation model must, therefore, provide two things: a way of choosing an ancestral allele for each new variant site, and a way of choosing a derived allele given the parental allele at each mutation. Perhaps the simplest mutation model implemented in `msprime` is the `InfiniteAlleles` mutation model, which keeps an internal counter so that the requested alleles are assigned subsequent (and therefore unique) integers.

The distribution of ancestral alleles is used to choose the allele present at the root of the tree at each mutated site, i.e., the `root_distribution`. Mutation models with a finite possible set of alleles have a natural choice for this distribution—the *stationary distribution* of the mutation process. (All mutation models are Markovian, so this may be found as the top left eigenvector of the mutation matrix.) This is the default in most models, except, e.g., the `BinaryMutationModel`, whose alleles are 0 and 1 and always labels the ancestral allele “0”. However, mutational processes are not in general stationary, so we often allow a different root distribution to be specified.

Since the general algorithm above applies mutations at a single rate independent of ancestral state, a model in which different alleles mutate at different rates must necessarily produce some silent mutations, i.e., mutations in which the derived allele is equal to the parental allele. To illustrate this, consider a mutation model in which *A* or *T* mutates to a randomly chosen different nucleotide at rate *α* and *C* or *G* mutates at rate *β*, with *β* < *α*. To implement this, first place mutations at the largest total rate, which is *α*. Then, at each site, choose an ancestral allele from the root distribution, and for each mutation, choose a derived allele as follows: if the parental allele is A or T, then choose a random derived allele different to the parental allele; if the parental allele is C or G, then choose the derived allele to be equal to the parent allele with probability *β*/(*α* + *β*), and randomly choose a different nucleotide otherwise. This produces the correct distribution by Poisson thinning: a Poisson process with rate *α* in which each point is discarded independently with probability *β*/(*α* + *β*) is equivalent to a Poisson process with rate *β*. All finite-state models (implemented under the generic `MatrixMutationModel` class) work in this way: mutations are placed at the maximum mutation rate, and then some silent mutations will result.

In previous versions of `msprime`, silent mutations were disallowed, and we could have removed them from the output entirely. However, we have chosen to leave them in, so that for instance simulating with the HKY mutation model will result in silent mutations if not all equilibrium frequencies are the same. The presence of silent mutations may at first be surprising but there is a good reason to leave them in: to allow layering of different mutation models. Suppose that we wanted to model the mutation process as a mixture of more than one model, e.g., Jukes-Cantor mutations at rate *μ*_1_, and HKY mutations occur at rate `μ`_2_. Layering multiple calls to `sim_mutations` is allowed, so we could first apply mutations with the `JC69` model at rate *μ*_1_ and then add more with the `HKY` model at rate *μ*_2_. However, there is a small statistical problem: suppose that after applying Jukes-Cantor mutations we have an A → C mutation, but then the HKY mutations inserts another mutation in the middle, resulting in A → C → C. If neither mutation model allows silent transitions, then this is clearly not correct, i.e., it is not equivalent to a model that simultaneously applies the two models. (The impact is small, however, as it only affects sites with more than one mutation.) The solution is to make the Jukes-Cantor model *state-independent* (also called “parent-independent”), by placing mutations at rate 4/3*μ*_1_ and then choosing the derived state for each mutation *independently* of the parent (so that 1/4 of mutations will be silent). If so—and, more generally, if the first mutational process put down is state-independent—then the result of sequentially applying the two mutation models is equivalent to the simultaneous model. To facilitate this, many mutation models have a `state_independent` option that increases the number of silent mutations and makes the model closer to state-independent.

Silent mutations are fully supported by `tskit`, which correctly accounts for their presence when computing statistics and performing other operations. For example, silent mutations have no effect on calculations of nucleotide site diversity.

### Time complexity of Hudson’s algorithm

As discussed in the Recombination section, the time complexity of Hudson’s algorithm is predicted to be quadratic in the population scaled recombination rate *ρ* = 4*N_e_L* (where *L* is the length of the genome in units of recombination distance) by Eq. (1). Fig. 8 shows the running time for simulations with a variety of population sizes, chromosome length and sample sizes, and shows this quadratic prediction is well supported by observations (see also Kelleher et al., 2016, Fig. 2). We also see that the dependence on *n* is quite weak, since increasing sample size 100-fold only increases run time by a factor of 2 or so. However, the log^2^ *n* factor implied by Eq. (1) (the sum is a harmonic number and can be approximated by log *n*) is not well supported by observed run times (or numbers of events) except possibly at very large values of *ρ*. It therefore appears that a different dependence on *n* is required to accurately predict simulation time for a given *ρ* and *n*.

**Figure 8:**
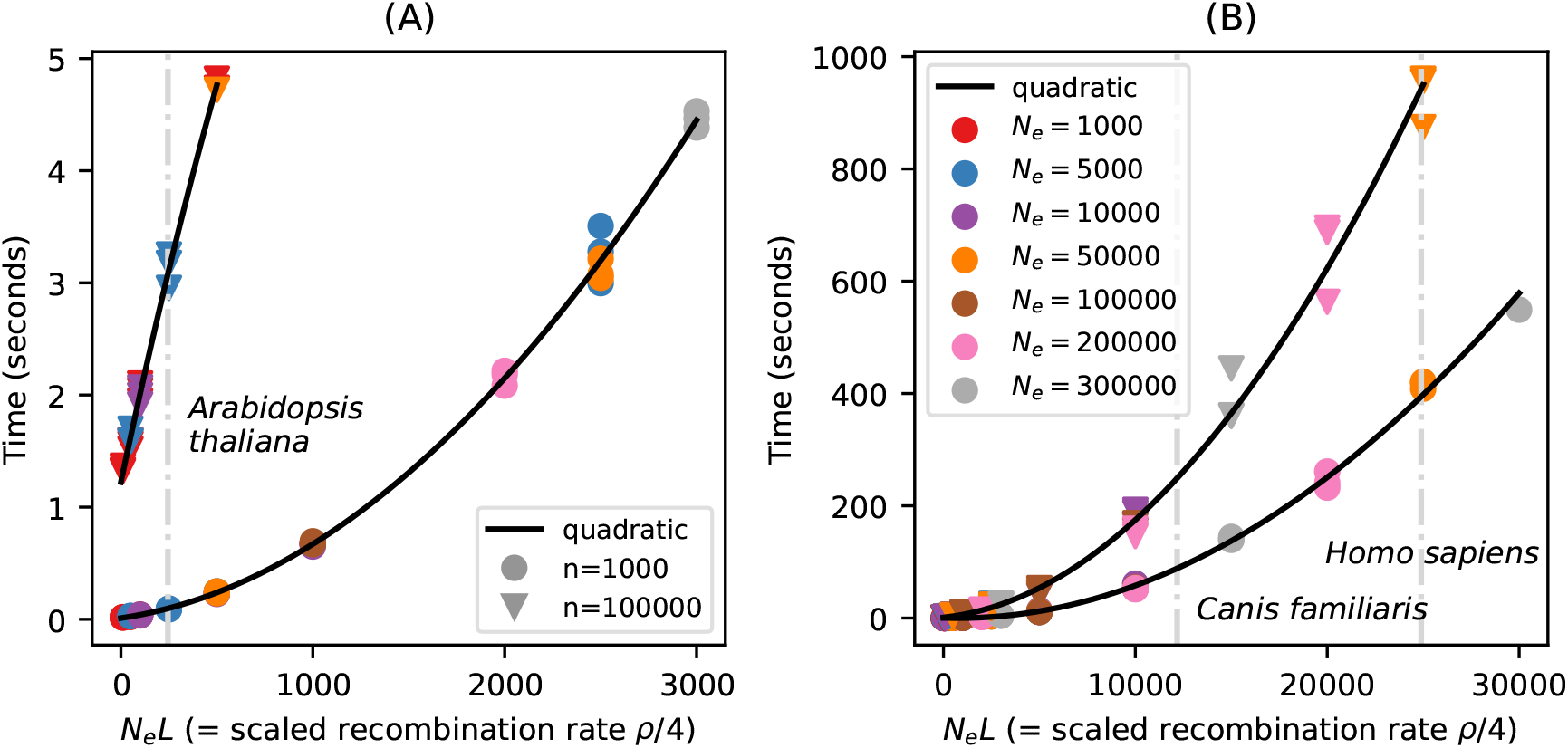
Running time of `sim_ancestry` for (A) small and (B) larger simulations on an Intel i7- 6600U CPU. Each point is the run time of one simulation, for various values of effective population size (*N_e_*), chromosome length in Morgans (*L*), and number of diploid samples (*n*). Run time scales quadratically with the product of *N_e_* and *L*, shown on the horizontal axis. For example, (A) shows that 1,000 samples of 1 Morgan-length chromosomes from a population of *N_e_* = 2,000 diploids would take about 2 seconds, and (equivalently) that the same number of 0.01 Morgan segments with *N_e_* = 200,000 would take the same time. Since recombination rate in these simulations was 10^-8^, L is the number of base pairs divided by 10^8^. The black lines are quadratic fits separately in each panel and sample size. Vertical grey lines show the approximate values of N_e_L for chromosome 1 in three species, using values from the `stdpopsim` catalogue (Adrion et al., 2020a).

Fig. 8 is a useful yardstick, allowing us to predict how long simulations should take for a wide range of species. Taking a typical chromosome to be 1 Morgan in length, these plots show, roughly, that simulating chromosome-length samples from a population of thousands of individuals takes seconds, while samples from a population of tens of thousands take minutes. Simulating whole chromosomes for many species is very fast, with 1000 samples of chromosome 1 for *Arabidopsis thaliana* taking less than a second, and a few minutes for dogs and humans. However, the dependence on *ρ is* quadratic, and if *ρ* is sufficiently large simulations may not be feasible. For example, the *Drosophila melanogaster* chromosome 2L is about 23.5Mb long with an average recombination rate of around 2.4 × 10^-8^, so *L* ≈ 0.57, and with *N_e_* = 1.7 × 10^6^ (Li and Stephan, 2006), *N_e_L* ≈ 10^6^, so extrapolating the curve in Fig. 8B predicts that simulation would require around 177 hours for 1000 samples. For such large values of *ρ* we recommend users consider approximate simulations. Since `msprime` does not currently have efficient implementations of approximate coalescent with recombination models, in these cases we recommend using SMC based methods such as `scrm`, particularly if sample sizes are small. In practice, to predict the running time of a given simulation in `msprime`, we recommend that users measure run time in a series of simulations with short genome lengths and the desired sample size, and then predict run time by fitting a quadratic curve to genome length as in Fig. 8. It is important to note that the quadratic curves in the two panels of Fig. 8 are different, and predicting the run times of days-long simulations using the timing of seconds-long runs is unlikely to be very accurate. 1

What about simulations with changing population size? To understand how run time depends on demography it helps to consider why run time is quadratic in *ρ*. At any point in time, `msprime` must keep track of some number of lineages, each of which contains some number of chunks of genetic material. Common ancestor events reduce the number of lineages, and recombination events increase their number. However, with long genomes, only a small fraction of the common ancestor events will involve overlapping segments of ancestry and lead to coalescence in the marginal trees. Such disjoint segments are often far apart (on average, about distance *L*/2), and so recombine apart again immediately; it is these large numbers of rapid and inconsequential events that lead to the quadratic run time. The maximum number of lineages occurs when the increase and decrease in numbers of lineages due to common ancestor and recombination events balance out. To get an idea of run time we can estimate when this balance occurs. Suppose that the maximum number of lineages is *M*; at this time the rate of common ancestor events is *M*(*M* – 1)/(4*N_e_*) and the total rate of recombination is *Mℓ*, where *ℓ* is the mean length of genome carried by each lineage (including “trapped” non-ancestral material). At the maximum, coalescence and recombination rates are equal, so a typical segment of ancestry will spend roughly half its time in a lineage with at least one other such segment—and, since such lineages carry at least two segments, at most one-third of the lineages carry long trapped segments of ancestry. Since the maximum number of lineages is reached very quickly (Nelson et al., 2020), this implies that *ℓ* ≈ *L*/6. Setting the rates of recombination and common ancestor events to be equal and solving for *M*, we find that *M* is roughly equal to L*N_e_*. The number of lineages then decreases gradually from this maximum on the coalescent time scale, and therefore over roughly 2*N_e_* generations. Since the total rate of events when the maximum number of lineages is present is roughly *L*^2^*N_e_*/6, then the total number of events is proportional to (*LN_e_*)^2^—i.e., proportional to *ρ*^2^.

What does this tell us about run time for simulating time-varying population sizes? Suppose that population size today is *N*_1_, while *T* generations ago it was *N*_2_. Does the run time depend more on 4*N*_1_*L* or 4*N*_2_*L*? The answer depends on how *T* compares to *N*_1_: if *T/N*_1_ ≪ then the number of extant lineages remaining after *T* generations is likely to be substantial, and the algorithm runtime is primarily determined by *N*_2_. Conversely, if *T/N*_1_ ≫ 1, then few extant lineages are likely to remain by time *T* and runtime depends mainly on *N*_1_. For instance, in many agricultural species *N*_1_ ≈ 100, while *N*_2_ ≈ 10^5^, and the run time will depend critically on T—in other words, simulation will be quick in a species with a strong domestication bottleneck, and slow otherwise. 1

### Selective sweeps model

Sweep trajectories are generated in `msprime` using a jump process approximation to the conditional diffusion of an allele bound for fixation (Coop and Griffiths, 2004). The jump process moves back in time following the beneficial allele frequency, *p*, from some initial frequency (e.g., *p* =1) back to the origination of the allele at *p* = 1/(2*N*), tracking time in small increments *δt*. Then, given the frequency *p* at time *t*, the frequency *p*′ at time *t* + *δt* is given by

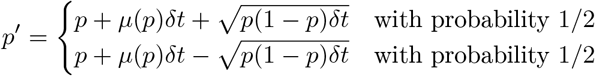

where

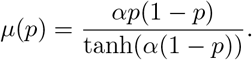

Here, *α* = 2*N s* and *s* is the fitness advantage in homozygotes. This model assumes genic selection (i.e., that the dominance coefficient *h* = 0.5), but can be generalised straightforwardly to include arbitrary dominance. We can also define trajectories to model neutral alleles and soft selective sweeps, which we plan as future additions to `msprime`.

### Likelihood calculations

We provide two functions to facilitate likelihood-based inference. Both are implemented only for the simplest case of the standard ARG with a constant population size, and require tree sequences compatible with the `record_full_arg` option as their arguments.

The `msprime.log_arg_likelihood`(`ts, r, N`) function returns the natural logarithm of the sampling probability of the tree sequence ts under the ARG with per-link, per-generation recombination probability `r` and population size `N` (e.g. Kuhner et al., 2000, equation (1)). Specifically, the function returns the logarithm of

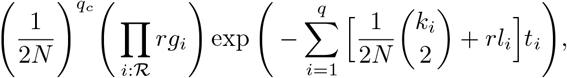

where *t_i_* is the number of generations between the (*i* – 1)th and *i*th event, *k_i_* is the number of extant ancestors in that interval, is the number of links in that interval that would split ancestral material should they recombine, *q* is the total number of events in the tree sequence *ts*, *q_c_* is the number of coalescences, 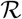 is the set of indices of time intervals which end in a recombination, and *g_i_* is the corresponding *gap*: the length of contiguous non-ancestral material around the link at which the recombination in question took place. The gap indicates the number of links (or length of genome in a continuous model) at which a recombination would result in exactly the observed pattern of ancestral material in the ARG. For a continuous model of the genome and a recombination in ancestral material, we set *g_i_* = 1 and interpret the result as a density.

The `msprime.unnormalised_log_mutation_likelihood`(`ts, m`) function returns the natural logarithm of the probability of the mutations recorded in the tree sequence ts given the corresponding ancestry, assuming the infinite sites model, up to a normalising constant which depends on the pattern of mutations, but not on the tree sequence or the per-site, per-generation mutation probability `m`. Specifically, the function returns the logarithm of

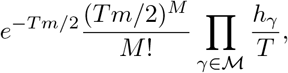

where *T* and 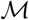 are the total branch length and set of mutations in `ts`, respectively, and for a mutation *γ, h_γ_* is the total branch length on which *γ* could have arisen while appearing on all of the leaves of ts it does, and on no others. Unary nodes on marginal trees arising from the `record_full_arg` option mean that, in general *h_γ_* corresponds to the length of one or more edges.

### Multiple merger coalescent model

Multiple merger coalescents, in which no more than one group of a random number of ancestral lineages may merge into a common ancestor at a given time, are referred to as Λ-coalescents. The rate at which a given group of *k* out of a total of *b* lineages merges is

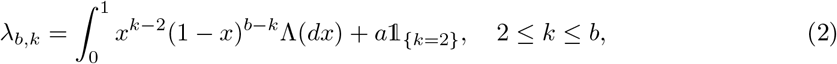

where 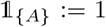 A holds, and zero otherwise, *a* ≦ 0 is a constant, and Λ is a finite measure on the unit interval without an atom at zero (Donnelly and Kurtz, 1999; Pitman, 1999; Sagitov, 1999). There is also a larger class of simultaneous multiple merger coalescents involving simultaneous mergers of distinct groups of lineages into several common ancestors (Schweinsberg, 2000). These are commonly referred to as Ξ-coalescents, and often arise from population models incorporating diploidy or more general polyploidy (Birkner et al., 2013a; Blath et al., 2016). To describe a general Ξ-coalescent, let △ denote the infinite simplex

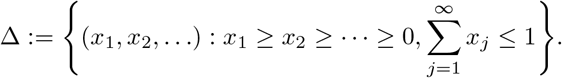

The rate of mergers is determined by Ξ = Ξ_0_ + *a*δ_0_, where *a* ≥ 0 is a constant, δo is the Dirac delta measure, and Ξ_0_ is a finite measure on Δ with no atom at (0, 0, …). For an initial number of blocks *b* ≥ 2 and *r* ∈ {1, 2,…, *b* – 1}, let *k*_1_ ≥ 2,…, *k_r_* ≥ 2 be the sizes of *r* merger events and *s* = *b* – *k*_1_ – · · · – *k_r_* be the number of blocks not participating in any merger. The rate of each possible set of mergers with sizes (*k*_1_,…, *k_r_*) is

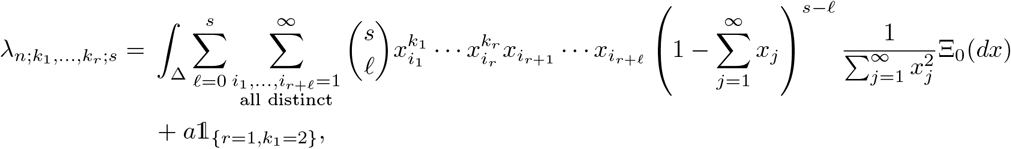

and the number of such (*k*_1_,…, *k_r_*) mergers is

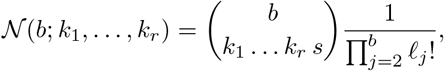

where 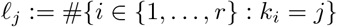 is the number of mergers of size *j* ≥ 2 (Schweinsberg, 2000).

Viewing coalescent processes strictly as mathematical objects, it is clear that the class of Ξ-coalescents contains Λ-coalescents as a specific example in which at most one group of lineages can merge at each time, and the class of Λ-coalescents contain the Kingman-coalescent as a special case. However, viewed as limits of ancestral processes derived from specific population models they are not nested. For example, one can obtain Λ-coalescents from haploid population models incorporating sweepstakes reproduction and high fecundity, and Ξ-coalescents for the same models for diploid populations (Birkner et al., 2013a). One should therefore apply the models as appropriate, i.e. Λ-coalescents to haploid (e.g. mtDNA) data, and Ξ-coalescents to diploid or polyploid (e.g. autosomal) data (Blath et al., 2016).

In `msprime` we have incorporated two examples of multiple-merger coalescents. One is a diploid extension (Birkner et al., 2013a) of the haploid Moran model adapted to sweepstakes reproduction considered by Eldon and Wakeley (2006). Let *N* denote the population size, and take *ψ* ∈ (0,1] to be fixed. In every generation, with probability 1 – *ε_N_* a single individual (picked uniformly at random) perishes. With probability *ε_N_*, ⌊*ψN*⌋ individuals picked uniformly without replacement perish instead. In either case, a surviving individual picked uniformly at random produces enough offspring to restore the population size back to *N*. Taking *ε_N_* = 1/*N^γ^* for some *γ* > 0, Eldon and Wakeley (2006) obtain Λ-coalescents for which the *Λ* measure in (2) is a point mass at *ψ*. The simplicity of this model does allow one to obtain some explicit mathematical results (see e.g. Der et al. (2012); Eldon and Freund (2018); Freund (2020); Matuszewski et al. (2018)), and the model has also been used to simulate gene genealogies within phylogenies (Zhu et al., 2015). As well as the haploid model of Eldon and Wakeley (2006), `msprime` provides the diploid version of Birkner et al. (2013a), in which individuals perish as above, but replacements are generated by sampling a single pair of diploid individuals as parents, with children sampling one chromosome from each parent. Hence, there are four parent chromosomes involved in each reproduction event, which can lead to up to four simultaneous mergers, giving rise to a Ξ-coalescent with merger rate

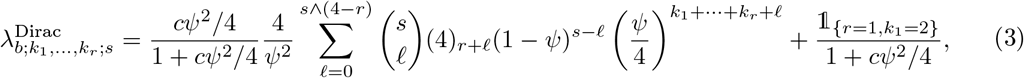

The interpretation of (3) is that ‘small’ reproduction events in which two lineages merge occur at rate 1/(1 + *cψ*^2^/4), while large reproduction events with the potential to result in simultaneous multiple mergers occur at rate (*cψ*^2^/4)/(1 + *cψ*^2^/4).

The other multiple merger coalescent model incorporated in `msprime` is the haploid population model considered by Schweinsberg (2003), as well as its diploid extension (Birkner et al., 2018). In the haploid version, in each generation of fixed size *N*, individuals produce random numbers of juveniles (*X*_1_,…, *X_N_*) independently, each distributed according to a stable law satisfying

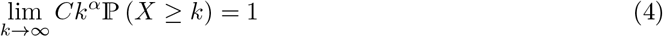

with index *α* > 0, and where *C* > 0 is a normalising constant. If the total number of juveniles *S_N_* := *X*_1_ +… + *X_N_* produced in this way is at least *N*, then *N* juveniles are sampled uniformly at random without replacement to form the next generation. As long as 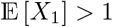, one can show that {*S_N_* < *N*} has exponentially small probability in *N*, and does not affect the resulting coalescent as *N* → ∞ (Schweinsberg, 2003). If *α* ≥ 2 the ancestral process converges to the Kingman-coalescent; if 1 ≤ *α* < 2 the ancestral process converges to a Λ-coalescent with Λ in (2) given by the Beta(2 – *α, α*) distribution, i.e.

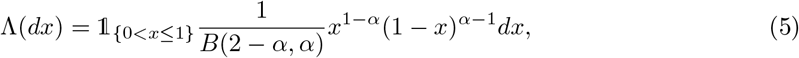

where *B*(*a, b*) = Γ(*a*)Γ(*b*)/Γ(*a*+*b*) for *α, b* > 0 is the beta function (Schweinsberg, 2003). This model has been adapted to diploid populations by Birkner et al. (2018), and the resulting coalescent is Ξ-coalescent with merger rate

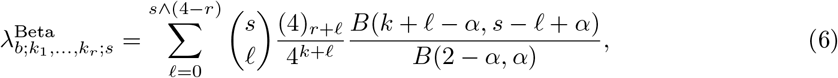

where *k* := *k*_1_ + … + *k_r_* (Blath et al., 2016; Birkner et al., 2018). The interpretation of (6) is that the random number of lineages participating in a potential merger is governed by the Λ-coalescent with rate (5), and all participating lineages are randomly allocated into one of four groups corresponding to the four parental chromosomes, giving rise to up to four simultaneous mergers.

The stable law (4) assumes that individuals can produce arbitrarily large numbers of juveniles. Since juveniles are at least fertilised eggs, it may be desirable to suppose that the number of juveniles surviving to reproductive maturity cannot be arbitrarily large. Hence we also consider an adaptation of the Schweinsberg (2003) model, where the random number of juveniles has a deterministic upper bound *ϕ*(*N*), and the distribution of the number of juveniles produced by a given parent (or pair of parents in the diploid case) is

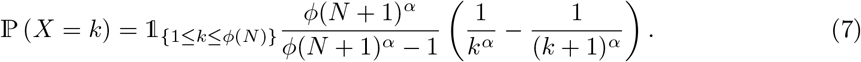

See Eldon and Stephan (2018) for a related model. One can follow the calculations of Schweinsberg (2003) or Birkner et al. (2018) to show that if 1 < *α* < 2 then, recalling that *k* = *k*_1_ + · · · + *k_r_*, the merger rate is

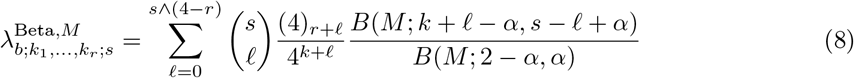

where 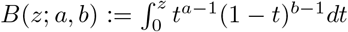 for *a,b* > 0 and 0 < *z* ≤ 1 is the incomplete beta function, and

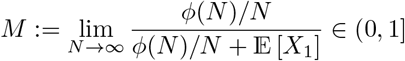

(Chetwynd-Diggle et al., 2022). In other words, the measure Λ driving the multiple mergers is of the same form as in (5) with 0 < *x* ≤ *M* in the case 1 < *α* < 2 and lim_*N*→∞_ *ϕ*(*N*)/*N* > 0. If *α* ≥ 2 or *ϕ*(*N*)/*N* → 0 then the ancestral process converges to the Kingman-coalescent (Chetwynd-Diggle et al., 2022).

## References

Jeffrey R Adrion, Christopher B Cole, Noah Dukler, Jared G Galloway, Ariella L Gladstein, Graham Gower, Christopher C Kyriazis, Aaron P Ragsdale, Georgia Tsambos, Franz Baumdicker, et al. A community-maintained standard library of population genetic models. Elife, 9:e54967, 2020a.

Jeffrey R Adrion, Jared G Galloway, and Andrew D Kern. Predicting the landscape of recombination using deep learning. Molecular biology and evolution, 37(6):1790–1808, 2020b.

Miguel Arenas. Simulation of molecular data under diverse evolutionary scenarios. PLoS Computational Biology, 8(5):e1002495, 2012.

Miguel Arenas and David Posada. Recodon: coalescent simulation of coding DNA sequences with recombination, migration and demography. BMC bioinformatics, 8(1):1–4, 2007.

Einar Árnason. Mitochondrial cytochrome *b* DNA variation in the high-fecundity Atlantic cod: trans-Atlantic clines and shallow gene genealogy. Genetics, 166(4):1871–1885, 2004.

Nicholas H Barton, Jerome Kelleher, and Alison M Etheridge. A new model for extinction and recolonization in two dimensions: quantifying phylogeography. Evolution: International journal of organic evolution, 64(9):2701–2715, 2010.

Franz Baumdicker and Peter Pfaffelhuber. The infinitely many genes model with horizontal gene transfer. Electronic Journal of Probability, 19:1–27, 2014. doi: 10.1214/EJP.v19-2642.

Mark A Beaumont, Wenyang Zhang, and David J Balding. Approximate Bayesian computation in population genetics. Genetics, 162(4):2025–2035, 2002.

Arnaud Becheler and L Lacey Knowles. Occupancy spectrum distribution: application for coalescence simulation with generic mergers. Bioinformatics, 02 2020. ISSN 1367-4803. btaa090.

Arnaud Becheler, Camille Coron, and Stéphane Dupas. The quetzal coalescence template library: A C++ programmers resource for integrating distributional, demographic and coalescent models. Molecular ecology resources, 19(3):788–793, 2019.

Andrew T Beckenbach. Mitochondrial haplotype frequencies in oysters: neutral alternatives to selection models. In Non-neutral evolution, pages 188–198. Springer, 1994.

Anand Bhaskar, Andrew G Clark, and Yun S Song. Distortion of genealogical properties when the sample is very large. Proceedings of the National Academy of Sciences, 111(6):2385–2390, 2014.

Matthias Birkner, Jochen Blath, Martin Möhle, Matthias Steinrücken, and Johanna Tams. A modified lookdown construction for the xi-fleming-viot process with mutation and populations with recurrent bottlenecks. Alea, 6:25–61, 2009.

Matthias Birkner, Jochen Blath, and Bjarki Eldon. An ancestral recombination graph for diploid populations with skewed offspring distribution. Genetics, 193(1):255–290, 2013a.

Matthias Birkner, Jochen Blath, and Bjarki Eldon. Statistical properties of the site-frequency spectrum associated with Λ-coalescents. Genetics, 195(3):1037–1053, 2013b.

Matthias Birkner, Huili Liu, and Anja Sturm. Coalescent results for diploid exchangeable population models. Electronic Journal of Probability, 23:1–44, 2018.

Jochen Blath, Mathias Christensen Cronjäger, Bjarki Eldon, and Matthias Hammer. The site-frequency spectrum associated with Ξ-coalescents. Theoretical Population Biology, 110:36–50, 2016.

Michael G.B. Blum and Olivier François. Non-linear regression models for Approximate Bayesian Computation. Statistics and Computing, 20(1):63–73, 2010.

Kevin S Bonham and Melanie I Stefan. Women are underrepresented in computational biology: An analysis of the scholarly literature in biology, computer science and computational biology. PLoS computational biology, 13(10):e1005134, 2017.

John M Braverman, Richard R Hudson, Norman L Kaplan, Charles H Langley, and Wolfgang Stephan. The hitchhiking effect on the site frequency spectrum of DNA polymorphisms. Genetics, 140(2):783–796, 1995.

Thomas Brown, Xavier Didelot, Daniel J. Wilson, and Nicola De Maio. SimBac: simulation of whole bacterial genomes with homologous recombination. Microbial Genomics, 2(1):1–6, 2016.

Lynsey Bunnefeld, Laurent A. F. Frantz, and Konrad Lohse. Inferring bottlenecks from genome-wide samples of short sequence blocks. Genetics, 201(3):1157–1169, 2015. doi: 10.1534/genet-ics.115.179861.

Clare Bycroft, Colin Freeman, Desislava Petkova, Gavin Band, Lloyd T Elliott, Kevin Sharp, Allan Motyer, Damjan Vukcevic, Olivier Delaneau, Jared O’Connell, et al. The UK Biobank resource with deep phenotyping and genomic data. Nature, 562:203–209, 2018.

Reed A Cartwright. DNA assembly with gaps (Dawg): simulating sequence evolution. Bioinformatics, 21(Suppl_3):iii31–iii38, 2005.

Antonio Carvajal-Rodríguez. Simulation of genomes: a review. Curr Genomics, 9(3):155, 2008.

Jeffrey Chan, Valerio Perrone, Jeffrey P Spence, Paul A Jenkins, Sara Mathieson, and Yun S Song. A likelihood-free inference framework for population genetic data using exchangeable neural networks. Advances in neural information processing systems, 31:8594, 2018.

Brian Charlesworth and Jeffrey D Jensen. Effects of selection at linked sites on patterns of genetic variability. Annual Review of Ecology, Evolution, and Systematics, 52:177–197, 2021.

Brian Charlesworth, MT Morgan, and Deborah Charlesworth. The effect of deleterious mutations on neutral molecular variation. Genetics, 134(4):1289–1303, 1993.

Deborah Charlesworth, Brian Charlesworth, and MT Morgan. The pattern of neutral molecular variation under the background selection model. Genetics, 141(4):1619–1632, 1995.

Gary K Chen, Paul Marjoram, and Jeffrey D Wall. Fast and flexible simulation of DNA sequence data. Genome research, 19(1):136–142, 2009.

Hua Chen and Kun Chen. Asymptotic distributions of coalescence times and ancestral lineage numbers for populations with temporally varying size. Genetics, 194(3):721–736, 2013.

Jian-Min Chen, David N Cooper, Nadia Chuzhanova, Claude Férec, and George P Patrinos. Gene conversion: mechanisms, evolution and human disease. Nature Reviews Genetics, 8(10):762–775, 2007.

Jonathan A Chetwynd-Diggle, Bjarki Eldon, and Alison M Etheridge. Beta-coalescents when sample size is large. in preparation, 2022.

Lounès Chikhi, Willy Rodríguez, Simona Grusea, Patricia Santos, Simon Boitard, and Olivier Mazet. The IICR (inverse instantaneous coalescence rate) as a summary of genomic diversity: insights into demographic inference and model choice. Heredity, 120(1):13–24, 2018.

Graham Coop and Robert C Griffiths. Ancestral inference on gene trees under selection. Theoretical population biology, 66(3):219–232, 2004.

Jean-Marie Cornuet, Filipe Santos, Mark A Beaumont, Christian P Robert, Jean-Michel Marin, David J Balding, Thomas Guillemaud, and Arnaud Estoup. Inferring population history with DIY ABC: a user-friendly approach to approximate Bayesian computation. Bioinformatics, 24 (23):2713–2719, 2008.

Katalin Csilléry, Michael GB Blum, Oscar E Gaggiotti, and Olivier François. Approximate Bayesian computation (ABC) in practice. Trends in ecology & evolution, 25(7):410–418, 2010.

Katalin Csilléry, Olivier François, and Michael G B Blum. abc: An R package for approximate Bayesian computation (ABC). Methods in Ecology and Evolution, 3(3):475–479, 2012.

M Dayhoff, R Schwartz, and B Orcutt. A model of evolutionary change in proteins. Atlas of protein sequence and structure, 5:345–352, 1978.

Nicola De Maio and Daniel J. Wilson. The bacterial sequential markov coalescent. Genetics, 206 (1):333–343, 2017.

Nicola De Maio, Lukas Weilguny, Conor R Walker, Yatish Turakhia, Russell Corbett-Detig, and Nick Goldman. phastsim: efficient simulation of sequence evolution for pandemic-scale datasets. bioRxiv, 2021.

Ricky Der, Charles Epstein, and Joshua B Plotkin. Dynamics of neutral and selected alleles when the offspring distribution is skewed. Genetics, 191(4):1331–1344, 2012.

Michael M Desai, Aleksandra M Walczak, and Daniel S Fisher. Genetic diversity and the structure of genealogies in rapidly adapting populations. Genetics, 193(2):565–585, 2013.

Peter Donnelly and Thomas G Kurtz. Particle representations for measure-valued population models. The Annals of Probability, 27(1):166–205, 1999.

Samantha Kristin Dung, Andrea López, Ezequiel Lopez Barragan, Rochelle-Jan Reyes, Ricky Thu, Edgar Castellanos, Francisca Catalan, Emilia Huerta-Sánchez, and Rori V Rohlfs. Illuminating women’s hidden contribution to historical theoretical population genetics. Genetics, 211(2):363–366, 2019.

Richard Durrett and Jason Schweinsberg. Approximating selective sweeps. Theoretical population biology, 66(2):129–138, 2004.

Bjarki Eldon and Fabian Freund. Genealogical properties of subsamples in highly fecund populations. Journal of Statistical Physics, 172(1):175–207, 2018.

Bjarki Eldon and Wolfgang Stephan. Evolution of highly fecund haploid populations. Theoretical population biology, 119:48–56, 2018.

Bjarki Eldon and John Wakeley. Coalescent processes when the distribution of offspring number among individuals is highly skewed. Genetics, 172(4):2621–2633, 2006.

SN Ethier and RC Griffiths. On the two-locus sampling distribution. Journal of Mathematical Biology, 29(2):131–159, 1990.

Gregory Ewing and Joachim Hermisson. MSMS: a coalescent simulation program including recombination, demographic structure, and selection at a single locus. Bioinformatics, 26(16): 2064–2065, 2010.

Laurent Excoffier and Matthieu Foll. Fastsimcoal: a continuous-time coalescent simulator of genomic diversity under arbitrarily complex evolutionary scenarios. Bioinformatics, 27(9):1332–1334, 2011.

Joseph Felsenstein and Gary A Churchill. A hidden markov model approach to variation among sites in rate of evolution. Molecular biology and evolution, 13(1):93–104, 1996.

Lex Flagel, Yaniv Brandvain, and Daniel R Schrider. The unreasonable effectiveness of convolutional neural networks in population genetic inference. Molecular biology and evolution, 36(2):220–238, 2019.

William Fletcher and Ziheng Yang. INDELible: a flexible simulator of biological sequence evolution. Molecular biology and evolution, 26(8):1879–1888, 2009.

Fabian Freund. Cannings models, population size changes and multiple-merger coalescents. Journal of mathematical biology, 80(5):1497–1521, 2020.

Nicolas Galtier, Frantz Depaulis, and N. H. Barton. Detecting bottlenecks and selective sweeps from DNA sequence polymorphism. Genetics, 155(2):981–987, 2000.

Paul P Gardner, James M Paterson, Stephanie R McGimpsey, Fatemeh Ashari Ghomi, Sinan U Umu, Aleksandra Pawlik, Alex Gavryushkin, and Michael A Black. Sustained software development, not number of citations or journal choice, is indicative of accurate bioinformatic software. bioRxiv, page 092205, 2021.

R. Chris Gaynor, Gregor Gorjanc, and John M. Hickey. AlphaSimR: An R-package for breeding program simulations. G3: Genes, Genomes, Genetics, 11, 2021. doi: 10.1093/g3journal/jkaa017.

John H Gillespie. Genetic drift in an infinite population: the pseudohitchhiking model. Genetics, 155(2):909–919, 2000.

Ariella L. Gladstein, Consuelo D Quinto-Cortés, Julian L. Pistorius, David Christy, Logan Gant-ner, and Blake L. Joyce. Simprily: A Python framework to simplify high-throughput genomic simulations. SoftwareX, 7:335–340, 2018.

Graham Gower, Aaron P Ragsdale, et al. Demes: a standard format for demographic models. In preparation, 2022.

Robert C Griffiths. The two-locus ancestral graph. Lecture Notes-Monograph Series, 18:100–117, 1991.

Robert C Griffiths and Paul Marjoram. An ancestral recombination graph. In P. Donnelly and S. Tavaré, editors, Progress in Population Genetics and Human Evolution, IMA Volumes in Mathematics and its Applications, volume 87, pages 257–270. Springer-Verlag, Berlin, 1997.

Robert C. Griffiths, Simon Tavare, Walter Fred Bodmer, and Peter James Donnelly. Sampling theory for neutral alleles in a varying environment. Philosophical Transactions of the Royal Society of London. Series B: Biological Sciences, 344(1310):403–410, 1994.

Frédéric Guillaume and Jacques Rougemont. Nemo: an evolutionary and population genetics programming framework. Bioinformatics, 22(20):2556–2557, 2006.

Benjamin C Haller and Philipp W Messer. SLiM 3: forward genetic simulations beyond the Wright–Fisher model. Molecular biology and evolution, 36(3):632–637, 2019.

Benjamin C Haller, Jared Galloway, Jerome Kelleher, Philipp W Messer, and Peter L Ralph. Tree-sequence recording in SLiM opens new horizons for forward-time simulation of whole genomes. Molecular ecology resources, 2018.

Charles R Harris, K Jarrod Millman, Stéfan J van der Walt, Ralf Gommers, Pauli Virtanen, David Cournapeau, Eric Wieser, Julian Taylor, Sebastian Berg, Nathaniel J Smith, et al. Array programming with numpy. Nature, 585(7825):357–362, 2020.

Kelley Harris. From a database of genomes to a forest of evolutionary trees. Nature genetics, 51 (9):1306–1307, 2019.

Dennis Hedgecock. Does variance in reproductive success limit effective population sizes of marine organisms? Genetics and evolution of aquatic organisms, pages 122–134, 1994.

Dennis Hedgecock and Alexander I Pudovkin. Sweepstakes reproductive success in highly fecund marine fish and shellfish: a review and commentary. Bulletin of Marine Science, 87(4):971–1002, 2011.

Jotun Hein, Mikkel Schierup, and Carsten Wiuf. Gene genealogies, variation and evolution: a primer in coalescent theory. Oxford University Press, USA, 2004.

Joseph Heled and Alexei J Drummond. Bayesian inference of species trees from multilocus data. Molecular biology and evolution, 27(3):570–580, 2009.

Garrett Hellenthal and Matthew Stephens. mshot: modifying Hudson’s ms simulator to incorporate crossover and gene conversion hotspots. Bioinformatics, 23(4):520–521, 2007.

Steven Henikoff and Jorja G Henikoff. Amino acid substitution matrices from protein blocks. Proceedings of the National Academy of Sciences, 89(22):10915–10919, 1992.

Michael J Hickerson, Eli Stahl, and Naoki Takebayashi. msBayes: pipeline for testing comparative phylogeographic histories using hierarchical approximate bayesian computation. BMC bioinformatics, 8(1):1–7, 2007.

Sean Hoban, Giorgio Bertorelle, and Oscar E Gaggiotti. Computer simulations: tools for population and evolutionary genetics. Nature Reviews Genetics, 13(2):110–122, 2012.

Asger Hobolth and Jens Ledet Jensen. Markovian approximation to the finite loci coalescent with recombination along multiple sequences. Theoretical population biology, 98:48–58, 2014.

Asger Hobolth, Arno Siri-Jegousse, and Mogens Bladt. Phase-type distributions in population genetics. Theoretical population biology, 127:16–32, 2019.

Wen Huang, Naoki Takebayashi, Yan Qi, and Michael J Hickerson. MTML-msBayes: approximate Bayesian comparative phylogeographic inference from multiple taxa and multiple loci with rate heterogeneity. BMC bioinformatics, 12(1):1–14, 2011.

Richard R. Hudson. Properties of a neutral allele model with intragenic recombination. Theoretical Population Biology, 23:183–201, 1983a.

Richard R. Hudson. Testing the constant-rate neutral allele model with protein sequence data. Evolution, 37(1):203–217, 1983b.

Richard R. Hudson. Gene genealogies and the coalescent process. Oxford Surveys in Evolutionary Biology, 7:1–44, 1990.

Richard R. Hudson. Generating samples under a Wright-Fisher neutral model of genetic variation. Bioinformatics, 18(2):337–338, 2002.

Kristen K Irwin, Stefan Laurent, Sebastian Matuszewski, Severine Vuilleumier, Louise Ormond, Hyunjin Shim, Claudia Bank, and Jeffrey D Jensen. On the importance of skewed offspring distributions and background selection in virus population genetics. Heredity, 117(6):393–399, 2016.

P Johri, CF Aquadro, M Beaumont, B Charlesworth, L Excoffier, A Eyre-Walker, PD Keightley, M Lynch, G McVean, BA Payseur, et al. Statistical inference in population genomics. 2021.

Thomas H Jukes, Charles R Cantor, et al. Evolution of protein molecules. Mammalian protein metabolism, 3:21–132, 1969.

Jack Kamm, Jonathan Terhorst, Richard Durbin, and Yun S Song. Efficiently inferring the demo-graphic history of many populations with allele count data. Journal of the American Statistical Association, 115(531):1472–1487, 2020.

Norman Kaplan and Richard R. Hudson. The use of sample genealogies for studying a selectively neutral *m*-loci model with recombination. Theoretical Population Biology, 28:382–396, 1985.

Norman L Kaplan, Richard R Hudson, and Charles H Langley. The “hitchhiking effect” revisited. Genetics, 123(4):887–899, 1989.

Konrad J Karczewski, Laurent C Francioli, Grace Tiao, Beryl B Cummings, Jessica Alföldi, Qingbo Wang, Ryan L Collins, Kristen M Laricchia, Andrea Ganna, Daniel P Birnbaum, et al. The mutational constraint spectrum quantified from variation in 141,456 humans. Nature, 581(7809): 434–443, 2020.

Peter D. Keightley and Adam Eyre-Walker. Joint inference of the distribution of fitness effects of deleterious mutations and population demography based on nucleotide polymorphism frequencies. Genetics, 177(4):2251–2261, 2007.

Jerome Kelleher and Konrad Lohse. Coalescent simulation with msprime. In Julien Y. Dutheil, editor, Statistical Population Genomics, pages 191–230. Springer US, New York, NY, 2020.

Jerome Kelleher, Nicholas H Barton, and Alison M Etheridge. Coalescent simulation in continuous space. Bioinformatics, 29(7):955–956, 2013.

Jerome Kelleher, Alison M Etheridge, and Nicholas H Barton. Coalescent simulation in continuous space: Algorithms for large neighbourhood size. Theoretical population biology, 95:13–23, 2014.

Jerome Kelleher, Alison M Etheridge, and Gilean McVean. Efficient coalescent simulation and genealogical analysis for large sample sizes. PLoS computational biology, 12(5):e1004842, 2016.

Jerome Kelleher, Kevin R. Thornton, Jaime Ashander, and Peter L. Ralph. Efficient pedigree recording for fast population genetics simulation. PLoS Computational Biology, 14(11):1–21, 11 2018.

Jerome Kelleher, Yan Wong, Anthony W. Wohns, Chaimaa Fadil, Patrick K. Albers, and Gil McVean. Inferring whole-genome histories in large population datasets. Nature Genetics, 51(9): 1330–1338, 2019.

Andrew D Kern and Daniel R Schrider. Discoal: flexible coalescent simulations with selection. Bioinformatics, 32(24):3839–3841, 2016.

Yuseob Kim and Wolfgang Stephan. Detecting a local signature of genetic hitchhiking along a recombining chromosome. Genetics, 160(2):765–777, 2002.

Motoo Kimura. A simple method for estimating evolutionary rates of base substitutions through comparative studies of nucleotide sequences. Journal of molecular evolution, 16(2):111–120, 1980.

Motoo Kimura. Estimation of evolutionary distances between homologous nucleotide sequences. Proceedings of the National Academy of Sciences, 78(1):454–458, 1981.

John F. C. Kingman. The coalescent. Stochastic processes and their applications, 13(3):235–248, 1982a.

John FC Kingman. On the genealogy of large populations. Journal of applied probability, 19(A): 27–43, 1982b.

Thomas Kluyver, Benjamin Ragan-Kelley, Fernando Pérez, Brian Granger, Matthias Bussonnier, Jonathan Frederic, Kyle Kelley, Jessica Hamrick, Jason Grout, Sylvain Corlay, Paul Ivanov, Damián Avila, Safia Abdalla, and Carol Willing. Jupyter notebooks – a publishing format for reproducible computational workflows. In F. Loizides and B. Schmidt, editors, Positioning and Power in Academic Publishing: Players, Agents and Agendas, pages 87 – 90. IOS Press, 2016.

Katharine L Korunes and Mohamed A F Noor. Gene conversion and linkage: effects on genome evolution and speciation. Molecular Ecology, 26(1):351–364, 2017.

Jere Koskela. Multi-locus data distinguishes between population growth and multiple merger coalescents. Statistical applications in genetics and molecular biology, 17(3), 2018.

Jere Koskela and Maite Wilke Berenguer. Robust model selection between population growth and multiple merger coalescents. Mathematical biosciences, 311:1–12, 2019.

Mary K Kuhner, Jon Yamato, and Joseph Felsenstein. Maximum likelihood estimation of recombination rates from population data. Genetics, 156(3):1393–1401, 2000.

Marguerite Lapierre, Camille Blin, Amaury Lambert, Guillaume Achaz, and Eduardo P. C. Rocha. The impact of selection, gene conversion, and biased sampling on the assessment of microbial demography. Molecular Biology and Evolution, 33(7):1711–1725, 2016.

Haipeng Li and Wolfgang Stephan. Inferring the demographic history and rate of adaptive substitution in Drosophila. PLOS Genetics, 2(10):1–10, 10 2006.

Heng Li and Richard Durbin. Inference of human population history from individual whole-genome sequences. Nature, 475:493–496, 2011.

Youfang Liu, Georgios Athanasiadis, and Michael E Weale. A survey of genetic simulation software for population and epidemiological studies. Human genomics, 3(1):79, 2008.

Joao S Lopes, David Balding, and Mark A Beaumont. Popabc: a program to infer historical demographic parameters. Bioinformatics, 25(20):2747–2749, 2009.

Thomas Mailund, Mikkel H Schierup, Christian NS Pedersen, Peter JM Mechlenborg, Jesper N Madsen, and Leif Schauser. CoaSim: a flexible environment for simulating genetic data under coalescent models. BMC bioinformatics, 6(1):1–6, 2005.

Paul Marjoram and Jeff D Wall. Fast “coalescent” simulation. BMC Genet, 7:16, 2006.

Gabor T. Marth, Eva Czabarka, Janos Murvai, and Stephen T. Sherry. The allele frequency spectrum in genome-wide human variation data reveals signals of differential demographic history in three large world populations. Genetics, 166(1):351–372, 2004.

Alicia R Martin, Christopher R Gignoux, Raymond K Walters, Genevieve L Wojcik, Benjamin M Neale, Simon Gravel, Mark J Daly, Carlos D Bustamante, and Eimear E Kenny. Human demo-graphic history impacts genetic risk prediction across diverse populations. The American Journal of Human Genetics, 100(4):635–649, 2017.

Alicia R Martin, Christopher R Gignoux, Raymond K Walters, Genevieve L Wojcik, Benjamin M Neale, Simon Gravel, Mark J Daly, Carlos D Bustamante, and Eimear E Kenny. Erratum: Human demographic history impacts genetic risk prediction across diverse populations (the american journal of human genetics (2020) 107 (4)(583–588),(s000292972030286x),(10.1016/j. ajhg. 2020.08. 017)). American journal of human genetics, 107(4):788–789, 2020.

Iain Mathieson and Aylwyn Scally. What is ancestry? PLoS Genetics, 16(3):e1008624, 2020.

Sebastian Matuszewski, Marcel E Hildebrandt, Guillaume Achaz, and Jeffrey D Jensen. Coalescent processes with skewed offspring distributions and nonequilibrium demography. Genetics, 208(1): 323–338, 2018.

Jakob McBroome, Bryan Thornlow, Angie S Hinrichs, Nicola De Maio, Nick Goldman, David Haussler, Russell Corbett-Detig, and Yatish Turakhia. A daily-updated database and tools for comprehensive SARS-CoV-2 mutation-annotated trees. bioRxiv, 2021.

James R McGill, Elizabeth A Walkup, and Mary K Kuhner. GraphML specializations to codify ancestral recombinant graphs. Fron Genet, 4:146, 2013.

Patrick F McKenzie and Deren AR Eaton. ipcoal: An interactive Python package for simulating and analyzing genealogies and sequences on a species tree or network. Bioinformatics, 36(14): 4193–4196, 2020.

Gilean A. T. McVean and Niall J. Cardin. Approximating the coalescent with recombination. Philos Trans R Soc Lond B Biol Sci, 360:1387–1393, 2005.

Karen H Miga, Sergey Koren, Arang Rhie, Mitchell R Vollger, Ariel Gershman, Andrey Bzikadze, Shelise Brooks, Edmund Howe, David Porubsky, Glennis A Logsdon, et al. Telomere-to-telomere assembly of a complete human X chromosome. Nature, 585(7823):79–84, 2020.

Mark J Minichiello and Richard Durbin. Mapping trait loci by use of inferred ancestral recombination graphs. The American Journal of Human Genetics, 79(5):910–922, 2006.

Martin Möhle and Serik Sagitov. A classification of coalescent processes for haploid exchangeable population models. Annals of Probability, pages 1547–1562, 2001.

Francesco Montinaro, Vasili Pankratov, Burak Yelmen, Luca Pagani, and Mayukh Mondal. Revisiting the Out of Africa event with a novel deep learning approach. bioRxiv, 2020.

Richard A Neher and Oskar Hallatschek. Genealogies of rapidly adapting populations. Proceedings of the National Academy of Sciences, 110(2):437–442, 2013.

Dominic Nelson, Jerome Kelleher, Aaron P Ragsdale, Claudia Moreau, Gil McVean, and Simon Gravel. Accounting for long-range correlations in genome-wide simulations of large cohorts. PLoS genetics, 16(5):e1008619, 2020.

Rasmus Nielsen. Estimation of population parameters and recombination rates from single nucleotide polymorphism. Genetics, 154(2):931–942, 2000.

Matthew Osmond and Graham Coop. Estimating dispersal rates and locating genetic ancestors with genome-wide genealogies. bioRxiv, 2021.

Pier Francesco Palamara. ARGON: fast, whole-genome simulation of the discrete time Wright-Fisher process. Bioinformatics, 32(19):3032–3034, 2016.

Christian M Parobek, Frederick I Archer, Michelle E DePrenger-Levin, Sean M Hoban, Libby Liggins, and Allan E Strand. skelesim: an extensible, general framework for population genetic simulation in r. Molecular ecology resources, 17(1):101–109, 2017.

Pavlos Pavlidis, Stefan Laurent, and Wolfgang Stephan. msABC: a modification of Hudson’s ms to facilitate multi-locus ABC analysis. Molecular Ecology Resources, 10(4):723–727, 2010.

Stephan Peischl, E Koch, RF Guerrero, and Mark Kirkpatrick. A sequential coalescent algorithm for chromosomal inversions. Heredity, 111(3):200–209, 2013.

Bo Peng, Huann-Sheng Chen, Leah E Mechanic, Ben Racine, John Clarke, Elizabeth Gillanders, and Eric J Feuer. Genetic data simulators and their applications: an overview. Genetic epidemiology, 39(1):2–10, 2015.

Jim Pitman. Coalescents with multiple collisions. Annals of Probability, pages 1870–1902, 1999.

Pierre Pudlo, Jean Michel Marin, Arnaud Estoup, Jean Marie Cornuet, Mathieu Gautier, and Christian P. Robert. Reliable ABC model choice via random forests. Bioinformatics, 32(6): 859–866, 2016.

Consuelo D Quinto-Cortés, August E Woerner, Joseph C Watkins, and Michael F Hammer. Modeling SNP array ascertainment with Approximate Bayesian Computation for demographic inference. Scientific reports, 8(1):1–10, 2018.

Fernando Racimo, David Gokhman, Matteo Fumagalli, Amy Ko, Torben Hansen, Ida Moltke, Anders Albrechtsen, Liran Carmel, Emilia Huerta-Sánchez, and Rasmus Nielsen. Archaic adaptive introgression in TBX15/WARS2. Molecular Biology and Evolution, 34(3):509–524, 2017.

Aaron P Ragsdale, Dominic Nelson, Simon Gravel, and Jerome Kelleher. Lessons learned from bugs in models of human history. American Journal of Human Genetics, 107(4):583–588, 2020.

Peter Ralph, Kevin Thornton, and Jerome Kelleher. Efficiently summarizing relationships in large samples: a general duality between statistics of genealogies and genomes. Genetics, 215(3): 779–797, 2020.

Andrew Rambaut and Nicholas C Grassly. Seq-Gen: an application for the Monte Carlo simulation of DNA sequence evolution along phylogenetic trees. Bioinformatics, 13(3):235–238, 1997.

Matthew D Rasmussen, Melissa J Hubisz, Ilan Gronau, and Adam Siepel. Genome-wide inference of ancestral recombination graphs. PLoS genetics, 10(5):e1004342, 2014.

Louis Raynal, Jean Michel Marin, Pierre Pudlo, Mathieu Ribatet, Christian P. Robert, and Arnaud Estoup. ABC random forests for Bayesian parameter inference. Bioinformatics, 35(10):1720–1728, 2019.

Angel G Rivera-Colón, Nicolas C Rochette, and Julian M Catchen. Simulation with RADinitio improves RADseq experimental design and sheds light on sources of missing data. Molecular ecology resources, 21(2):363–378, 2021.

Benjamin K Rosenzweig, James B Pease, Nora J Besansky, and Matthew W Hahn. Powerful methods for detecting introgressed regions from population genomic data. Molecular ecology, 25 (11):2387–2397, 2016.

Serik Sagitov. The general coalescent with asynchronous mergers of ancestral lines. Journal of Applied Probability, 36(4):1116–1125, 1999.

Théophile Sanchez, Jean Cury, Guillaume Charpiat, and Flora Jay. Deep learning for population size history inference: Design, comparison and combination with approximate bayesian computation. Molecular Ecology Resources, 2020.

Nathan K Schaefer, Beth Shapiro, and Richard E Green. An ancestral recombination graph of human, Neanderthal, and Denisovan genomes. Science Advances, 7(29):eabc0776, 2021.

Stephan Schiffels and Richard Durbin. Inferring human population size and separation history from multiple genome sequences. Nat Genet, 46:919–925, 2014.

Daniel R Schrider and Andrew D Kern. Supervised machine learning for population genetics: a new paradigm. Trends in Genetics, 34(4):301–312, 2018.

Jason Schweinsberg. Coalescents with simultaneous multiple collisions. Electron Journal of Probability, 5:1–50, 2000.

Jason Schweinsberg. Coalescent processes obtained from supercritical Galton–Watson processes. Stochastic processes and their Applications, 106(1):107–139, 2003.

Jason Schweinsberg. Rigorous results for a population model with selection II: genealogy of the population. Electronic Journal of Probability, 22:1–54, 2017.

Geordan Shannon, Melanie Jansen, Kate Williams, Carlos Cáceres, Angelica Motta, Aloyce Odhi-ambo, Alie Eleveld, and Jenevieve Mannell. Gender equality in science, medicine, and global health: where are we at and why does it matter? The Lancet, 393(10171):560–569, 2019.

Sara Sheehan and Yun S Song. Deep learning for population genetic inference. PLoS computational biology, 12(3):e1004845, 2016.

Sara Sheehan, Kelley Harris, and Yun S Song. Estimating variable effective population sizes from multiple genomes: a sequentially markov conditional sampling distribution approach. Genetics, 194(3):647–662, 2013.

Ilya Shlyakhter, Pardis C. Sabeti, and Stephen F. Schaffner. Cosi2: an efficient simulator of exact and approximate coalescent with selection. Bioinformatics, 30(23):3427–3429, 2014.

Adam Siepel. Challenges in funding and developing genomic software: roots and remedies. Genome Biology, 20, 2019.

Leo Speidel, Marie Forest, Sinan Shi, and Simon R. Myers. A method for genome-wide genealogy estimation for thousands of samples. Nature Genetics, 51(9):1321–1329, 2019.

Leo Speidel, Lara Cassidy, Robert W Davies, Garrett Hellenthal, Pontus Skoglund, and Simon R Myers. Inferring population histories for ancient genomes using genome-wide genealogies. Molecular Biology and Evolution, 2021.

Jeffrey P Spence and Yun S Song. Inference and analysis of population-specific fine-scale recombination maps across 26 diverse human populations. Science Advances, 5(10):eaaw9206, 2019.

Chris CA Spencer and Graham Coop. SelSim: a program to simulate population genetic data with natural selection and recombination. Bioinformatics, 20(18):3673–3675, 2004.

Stephanie J Spielman and Claus O Wilke. Pyvolve: a flexible Python module for simulating sequences along phylogenies. PloS one, 10(9), 2015.

Paul R Staab and Dirk Metzler. Coala: an R framework for coalescent simulation. Bioinformatics, 32(12):1903–1904, 2016.

Paul R Staab, Sha Zhu, Dirk Metzler, and Gerton Lunter. scrm: Efficiently simulating long sequences using the approximated coalescent with recombination. Bioinformatics, 31(10):1680–1682, 2015.

Fumio Tajima. Evolutionary relationship of DNA sequences in finite populations. Genetics, 105(2):437–460, 1983. ISSN 0016-6731.

Lin Tang. Genealogy at the genome scale. Nature methods, 16(11):1077–1077, 2019.

Tomoya Tanjo, Yosuke Kawai, Katsushi Tokunaga, Osamu Ogasawara, and Masao Nagasaki. Prac-tical guide for managing large-scale human genome data in research. Journal of Human Genetics, 66(1):39–52, 2021.

Simon Tavaré et al. Some probabilistic and statistical problems in the analysis of DNA sequences. Lectures on mathematics in the life sciences, 17(2):57–86, 1986.

Drew E Terasaki Hart, Anusha P Bishop, and Ian J Wang. Geonomics: forward-time, spatially explicit, and arbitrarily complex landscape genomic simulations. Molecular Biology and Evolution, 2021.

Jonathan Terhorst, John A Kamm, and Yun S Song. Robust and scalable inference of population history from hundreds of unphased whole genomes. Nature genetics, 49(2):303–309, 2017.

Kosuke M Teshima and Hideki Innan. mbs: modifying Hudson’s ms software to generate samples of DNA sequences with a biallelic site under selection. BMC Bioinformatics, 10(1):166, 2009.

Kevin Thornton and Peter Andolfatto. Approximate Bayesian inference reveals evidence for a recent, severe bottleneck in a Netherlands population of Drosophila melanogaster. Genetics, 172 (3):1607–1619, 2006.

Kevin R Thornton. A C++ template library for efficient forward-time population genetic simulation of large populations. Genetics, 198(1):157–166, 2014.

Bianca Trinkenreich, Igor Wiese, Anita Sarma, Marco Gerosa, and Igor Steinmacher. Women’s par-ticipation in open source software: A survey of the literature. arXiv preprint arXiv:2105.08777, 2021.

Tskit developers. Tskit: a portable library for population scale genealogical analysis. In preparation, 2022.

Yatish Turakhia, Bryan Thornlow, Angie S Hinrichs, Nicola De Maio, Landen Gozashti, Robert Lanfear, David Haussler, and Russell Corbett-Detig. Ultrafast sample placement on existing trees (UShER) enables real-time phylogenetics for the SARS-CoV-2 pandemic. Nature Genetics, pages 1–8, 2021.

David LJ Vendrami, Lloyd S Peck, Melody S Clark, Bjarki Eldon, Michael Meredith, and Joseph I Hoffman. Sweepstake reproductive success and collective dispersal produce chaotic genetic patchiness in a broadcast spawner. Science advances, 7(37):eabj4713, 2021.

Thimothée Virgoulay, François Rousset, Camille Noûs, and Raphaël Leblois. Gspace: an exact coalescence simulator of recombining genomes under isolation by distance. Bioinformatics, 2021.

John Wakeley. Coalescent theory: an introduction. Roberts and Company, Englewood, Colorado, 2008.

John Wakeley, Léandra King, Bobbi S Low, and Sohini Ramachandran. Gene genealogies within a fixed pedigree, and the robustness of Kingman’s coalescent. Genetics, 190(4):1433–1445, 2012.

Ke Wang, Iain Mathieson, Jared O’Connell, and Stephan Schiffels. Tracking human population structure through time from whole genome sequences. PLoS Genetics, 16(3):e1008552, 2020.

Ying Wang and Bruce Rannala. Bayesian inference of fine-scale recombination rates using population genomic data. Philosophical Transactions of the Royal Society of London. Series B: Biological Sciences, 363(1512):3921–3930, 2008.

Ying Wang, Ying Zhou, Linfeng Li, Xian Chen, Yuting Liu, Zhi-Ming Ma, and Shuhua Xu. A new method for modeling coalescent processes with recombination. BMC Bioinformatics, 15(1):273, 2014.

Daniel Wegmann, Christoph Leuenberger, Samuel Neuenschwander, and Laurent Excoffier. ABC-toolbox: a versatile toolkit for approximate Bayesian computations. BMC bioinformatics, 11(1): 1–7, 2010.

Maren Wellenreuther and Sarah Otto. Women in evolution–highlighting the changing face of evolutionary biology. Evolutionary Applications, 9(1):3–16, 2016.

Peter R Wilton, Shai Carmi, and Asger Hobolth. The SMC’ is a highly accurate approximation to the ancestral recombination graph. Genetics, 200(1):343–355, 2015.

Carsten Wiuf and Jotun Hein. The ancestry of a sample of sequences subject to recombination. Genetics, 151(3):1217–1228, 1999a.

Carsten Wiuf and Jotun Hein. Recombination as a point process along sequences. Theoretical Population Biology, 55(3):248–259, 1999b.

Carsten Wiuf and Jotun Hein. The coalescent with gene conversion. Genetics, 155(1):451–462, 2000.

Anthony Wilder Wohns, Yan Wong, Ben Jeffery, Ali Akbari, Swapan Mallick, Ron Pinhasi, Nick Patterson, David Reich, Jerome Kelleher, and Gil McVean. A unified genealogy of modern and ancient genomes. bioRxiv, 2021.

Tao Yang, Hong-Wen Deng, and Tianhua Niu. Critical assessment of coalescent simulators in modeling recombination hotspots in genomic sequences. BMC Bioinformatics, 15:3, 2014.

Xiguo Yuan, David J Miller, Junying Zhang, David Herrington, and Yue Wang. An overview of population genetic data simulation. Journal of Computational Biology, 19(1):42–54, 2012.

Sha Zhu, James H Degnan, Sharyn J Goldstien, and Bjarki Eldon. Hybrid-Lambda: simulation of multiple merger and Kingman gene genealogies in species networks and species trees. BMC Bioinformatics, 16(292), 2015.

